# IntAct: a non-disruptive internal tagging strategy to study actin isoform organization and function

**DOI:** 10.1101/2021.10.25.465733

**Authors:** M.C. van Zwam, A. Dhar, W. Bosman, W. van Straaten, S. Weijers, E. Seta, B. Joosten, S. Palani, K. van den Dries

## Abstract

Actin plays a central role in many cell biological processes including division and motility. Mammals have six, highly conserved actin isoforms with nonredundant biological functions, yet the molecular basis of isoform specificity remains elusive due to a lack of tools. Here, we describe the development of IntAct, an internal tagging strategy to study actin isoform function in fixed and living cells. We first identified a residue pair in β-actin that permits non-disruptive tag integration. Next, we used knock-in cell lines to demonstrate that the expression and filament incorporation of IntAct β-actin is indistinguishable from wildtype. Furthermore, IntAct β-actin remains associated with actin-binding proteins profilin, cofilin and formin family members DIAPH1 and FMNL2 and can be targeted in living cells. To demonstrate the usability of IntAct for actin isoform investigations, we also generated IntAct γ-actin cells and show that actin isoform specific distribution remains unaltered in human cells. Moreover, introduction of tagged actin variants in yeast demonstrated an expected variant-dependent incorporation into patches and filaments. Together, our data indicate that IntAct is a versatile tool to study actin isoform localization, dynamics and molecular interactions.

## Introduction

Actin plays a central role during fundamental biological processes including cell division, shape maintenance, motility and contractility. In birds and mammals, actin has six isoforms, also called isoactins, which are encoded by different genes and expressed in a tissue and time-specific manner during development, homeostasis and pathology (1–3). All six isoactins have nonredundant functions as indicated by the discovery of disease-causing mutations in each of the genes encoding the isoforms (4). Although exceptions exist, it is generally acknowledged that four isoactins are expressed in muscle cells and two are ubiquitously expressed across tissues. The two ubiquitous isoactins, nonmuscle β- and γ-actin, display the highest similarity with only four different residues at their N-terminus (5). Despite this similarity, nonmuscle β- and γ-actin play specific roles in many actin-controlled processes including cell-cell junction formation (6), axon development (7), cell division (8, 9) and cell migration (10, 11). While it has been demonstrated that isoactin-specific posttranslational modification (12, 13), nucleation (8, 9) and translation speed (13, 14) contribute to the nonredundant role of β- and γ-actin in cellular processes, the molecular principles that govern the differential function of isoactins remain largely unclear. This is primarily due to the limited possibilities to specifically probe actin isoforms for biochemical and cell biological assays.

Common tools to label the actin cytoskeleton such as phalloidin (15) in fixed cells or Lifeact (16), F-tractin (17), UtrophinCH (18), and anti-actin nanobodies (19–21) in living cells do not discriminate between isoactins. Furthermore, C-terminal fusions of actin cannot be used to study isoactin biology since they only poorly assemble into filaments (22, 23). N-terminal fusions of actin have been used to study isoactin differences in cells (14, 24, 25), but the reporter tags are known to significantly interfere with actin dynamics (16), nucleation (26) and molecular interactions (16, 26–28). Moreover, N-terminal fusion prevents isoactin-specific and nonspecific posttranslational modifications crucial for proper actin function such as arginylation (12, 13) and acetylation (29, 30). Attempts to tag yeast actin internally with a tetra-cysteine tag for contractile ring visualization were unsuccessful since the modified actins were rejected by formins during filament elongation (31). An extended search for other internal sites that may be permissive for epitope tagging of actin has not been performed so far.

Here, we describe the development of a non-disruptive internal tagging strategy to study isoactin organization and function, which we call IntAct. For this, we first performed a microscopy-based screen for eleven internal actin positions and identified one residue pair that allows non-disruptive epitope tagging of actin. To prove its versatility and usability, we engineered CRISPR/Cas9-mediated knock-in cell lines with various antibody- and nanobody-based epitope tags in the identified position and demonstrate that the internally tagged actins are properly expressed and that the integration into filaments is unperturbed. By performing immunofluorescence, pulldown experiments and live-cell imaging with the internally tagged actins, we show that IntAct can be used to study isoactin localization, molecular interactions, and dynamics. Lastly, we show that internally tagged nonmuscle β- and γ- actin mimic the differential distribution of wildtype isoforms and demonstrate the incorporation of tagged actins variants into yeast actin patches and cables indicating the possibility to extend the use of IntAct to other species. Altogether, our results indicate that IntAct can provide unique insights into the isoactin-specific molecular principles that regulate cellular processes such as division, motility, and contractility.

## Results

### T229/A230 actin residue pair is permissive for epitope tag insertion

To identify a permissive residue pair to internally tag the actin protein, we first performed a medium-scale screen and tagged β-actin at eleven distinct residue pairs with a FLAG tag (**Fig. 1A**). These residue pairs were carefully selected, ensuring that at least one of the residues is part of an unstructured region (32) and that both residues were not involved in F-actin interactions (33). Furthermore, the first 40 residues were avoided since the coding mRNA for this region is involved in the different translation kinetics of actin isoforms (13). Eventually, two of the eleven selected positions were located in subdomain 3, seven in subdomain 4, and two were close to the ATP binding site (**Fig. S1**). C- and N-terminally tagged β-actin were included in the screen as a negative and positive control for filament integration, respectively. We chose the FLAG tag as an epitope for our screen because of its frequent use, small size (8 amino acids, DYKDDDDK), and availability of a highly specific and wellcharacterized antibody (34).

**Fig. 1.**
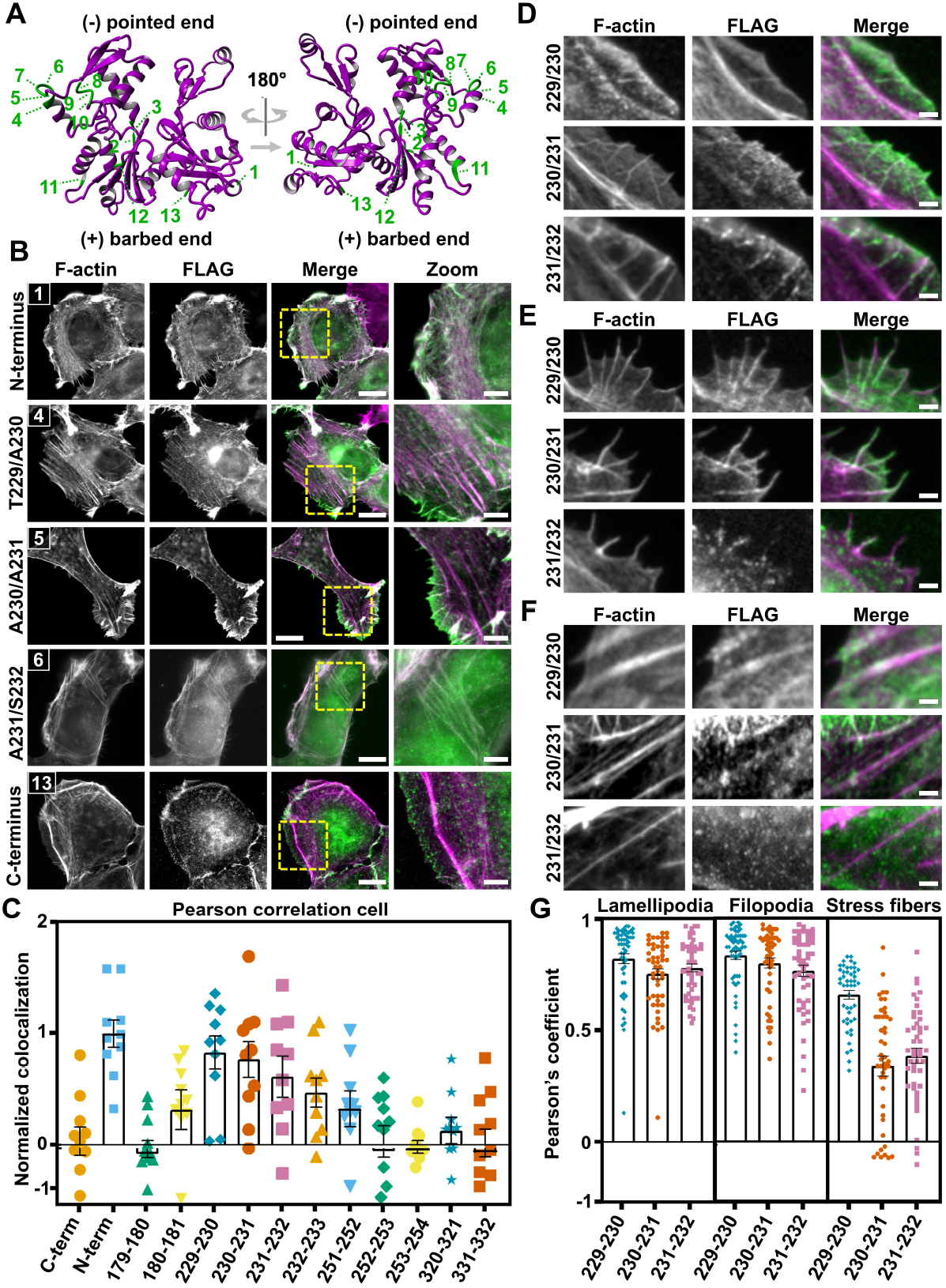
Identification of actin T229/A230 as a permissive target site for epitope tag integration. (**A**) Crystal structure of uncomplexed globular actin (magenta ribbon, PBD accession number: 1J6Z32) indicating the eleven internal target positions (green) as well as the N- and the C-terminus (1 and 13, respectively). (**B**) Representative widefield immunofluorescence images of F-actin (magenta) and FLAG (green) in HT1080 cells that overexpress the tagged β-actin variants. Shown are three internally tagged variants and the N- and C-terminally tagged β-actin. The remaining eight internally tagged variants are shown in **Suppl. Fig. S2**. Scale bar: 15 µm. Scale bar zoom: 5 µm. (**C**) Colocalization analysis of the microscopy results in **B** showing the normalized Pearson’s coefficient for each of the actin variants. Individual data points indicate single cells and in total, at least 10 cells from 2 independent experiments were included in the analysis. Bars represent the mean value, and error bars represent standard error of mean (SEM). (**D-F**) Representative images of (**D**) lamellipodia (**E**) filopodia and (**F**) stress fibers. Cells that overexpress the tagged β-actin variants were stained for F-actin (magenta) and FLAG (green). Scale bar: 2 µm. (**G**) Pearson’s colocalization analysis for the images in **D-F**. Individual data points indicate single lamellipodia, filopodia or stress fibers and in total, at least 40 structures in at least 20 different cells from 2 independent experiments were included in the analysis. Bars represent the mean value, and error bars represent standard error of mean (SEM).

To evaluate the integration of the tagged actins within the actin cytoskeleton, we overexpressed the thirteen actin variants in human fibrosarcoma cells (HT1080, **Fig. 1B-C, Fig. S2**) and retinal pigment epithelium cells (RPE1, **Fig. S3**) and performed an immunofluorescence staining for the FLAG tag and phalloidin as a total F-actin marker (**Fig. 1B, Fig. S2, Fig. S3A**). Interestingly, by visual inspection, we observed that most of the internally tagged actins were diffusely present within the cytosol with three notable exceptions (T229/A230, A230/A231 and A231/S232). Of these three variants, A230/A231 and A231/S232 only seemed to present a clear overlap with actin at the cell periphery but the T229/A230 overlapped almost entirely with the actin signal, similarly to N-terminally tagged actin. To quantify these observations, we first performed a Pearson correlation coefficient analysis on entire cells (**Fig. 1C, Fig. S3B**). As expected, N-terminally tagged actin showed a high degree of colocalization, while C-terminally tagged actin showed almost no colocalization. We therefore performed a unity-based normalization, adjusting the Pearson coefficient of the C- and N-terminus to zero and one, respectively, and normalized the other values within this window. While most of the internally tagged actin variants showed little to no colocalization, the T229/A230 variant displayed a high Pearson coefficient in both HT1080 and RPE1 cells (raw R^2^=0.68 and 0.75, respectively). Interestingly, for the A230/A231 and A231/S232 variant, we observed a very low colocalization in RPE1 cells, but a relatively high colocalization in HT1080 cells. We therefore sought to further dissect the integration of these three variants into the actin cytoskeleton of HT1080 cells and quantified the colocalization within individual actin-based structures (**Fig. 1D-F**). While we see a high degree of colocalization for all three variants in lamellipodia and filopodia, only the T229/A230 variant shows a high degree of colocalization also in stress fibers (**Fig. 1G**). This demonstrates the unique ability of the T229/A230 variant to be integrated in both branched as well as unbranched actin filament structures. Since we eventually aim to use our internal tagging strategy to study the molecular mechanisms of actin isoform diversity, we overexpressed internally tagged versions of the five other mammalian isoactins in HT1080 and RPE1 cells and observed that all isoforms were incorporated in various substructures of the actin cytoskeleton (**Fig. S4**). Together, these results strongly suggest that the T229/A230 residue pair in actin is permissive for epitope tag insertion.

### T229/A230 epitope tag insertion does not impair actin expression or assembly into filaments

To investigate the versatility and usability of the T229/A230 residue pair for epitope tag insertion, we applied CRISPR/Cas9-mediated homology-directed repair (HDR) to genetically introduce various tags into this position at the genomic locus of β-actin. We included the antibody-based epitope tags FLAG (DYKDDDDK), AU1 (DTYRYI) and AU5 (TDFYLK) as well as the recently developed nanobody(Nb)-based ALFA tag (PSRLEEELRRRLTEP) (35). Of note, we introduced the AU5 tag without the starting threonine (DFYLK), since this residue is redundant with the T229 of actin. Flow cytometry data shows that we retrieved a similar degree of knock-in efficiency for all the different tags (**Fig. S5A**). To specifically select for HDR events, we performed a ouabain-based coselection procedure (36) and again retrieved similar degree of knock-in efficiency for all the tags (**Fig. S5B**). For AU5, FLAG and ALFA, we continued to select clonal cell lines and for all three tags, based on western blot and sequencing analysis, we were able to retrieve clonal cell lines that exclusively produce internally tagged β-actin. For AU5 and FLAG, the clones were hemizygous and for ALFA, we obtained a homozygous knock-in clone (**Fig. S5C-E**). To also demonstrate that it is possible to tag multiple isoforms in the same cell, we performed a knock-in of the FLAG tag in γ-actin in the already established homozygous ALFA-β-actin HT1080 cells. Immunofluorescence labeling of FLAG and ALFA in these cells confirmed the possibility to simultaneously tag β- and γ-actin at the T229/A230 position with different epitope tags (**Fig. S6A**).

Previously published results indicated that a heterozygous knock-in of GFP into the genomic locus of β-actin for N-terminal tagging leads to a dramatic decrease of protein expression from the modified allele (27). After generation of our knock-in cell lines, we therefore first aimed to determine if internally tagging actin at position T229/A230 leads to an altered actin expression in the clonal cell lines. For this, we used heterozygous FLAG-β-actin cells since FLAG causes a gel shift on western blot, allowing a direct comparison of tagged and wildtype actin expression in the same cells. Quantification of the western blots demonstrated that the amount of FLAG-β-actin was similar to wildtype indicating that the cells did not downregulate the expression of β-actin from the knock-in allele (**Fig. 2A-B**). We also evaluated actin protein expression in the homozygous ALFA-β-actin cells and this showed that the total amount of actin was slightly lower in the ALFA-β-actin cells compared to parental HT1080 cells (**Fig. 2C-D**). Although we expect that this decrease is attributed to clonal variation, we wanted to exclude compromised global actin regulation in the ALFA-β-actin cells. We therefore evaluated the expression of γ-actin and α-smooth muscle actin (α-SMA) since a genetic loss of β-actin has been shown to induce the expression of these isoforms (37). Importantly, we neither observed differences in γ-actin expression or an induction of α-SMA in the ALFA-β-actin cells (**Fig. S6B-D**), strongly suggesting that global actin regulation is not perturbed by genetically tagging β-actin.

**Fig. 2.**
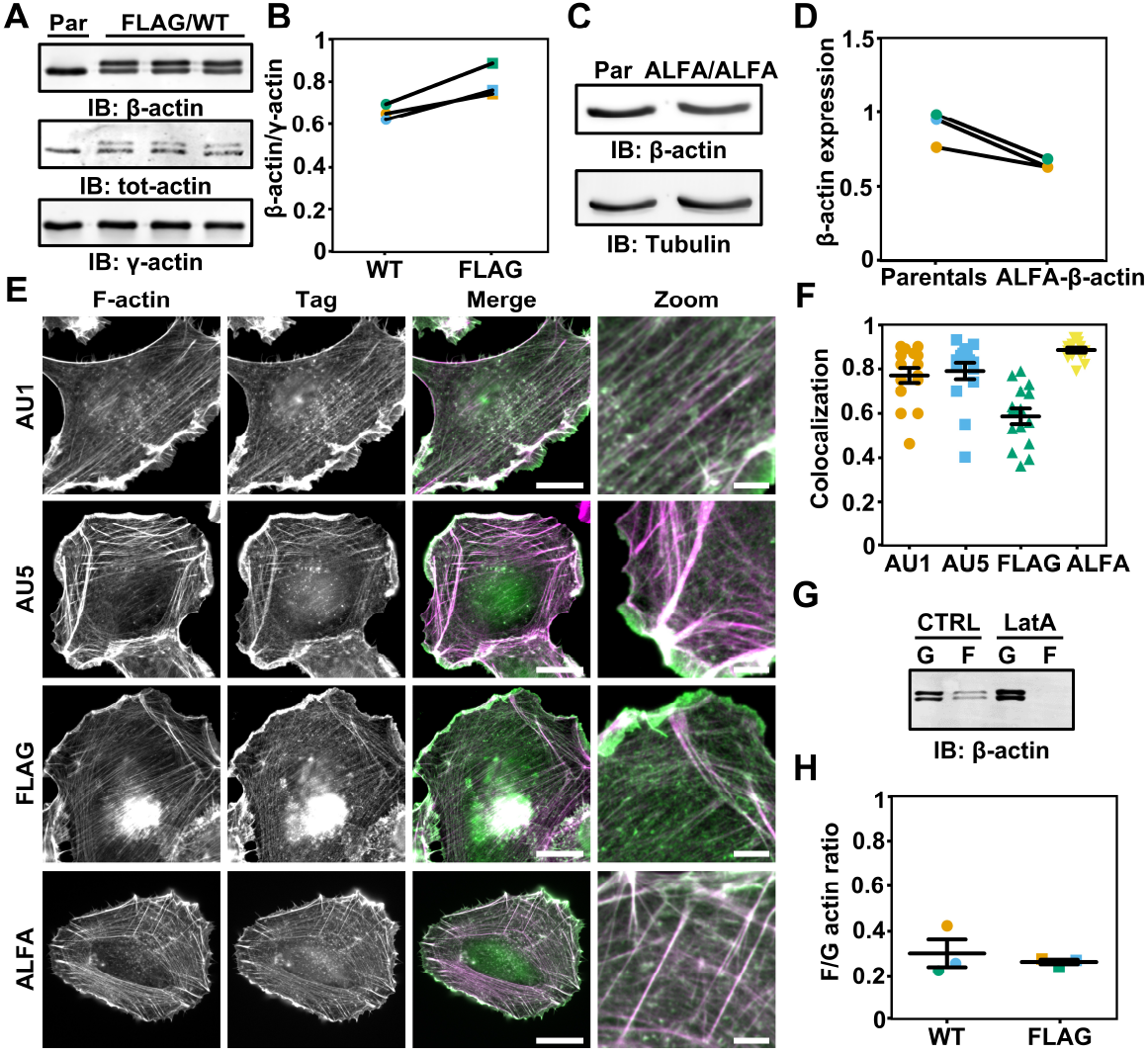
Actin functionality unperturbed by T229/A230 epitope integration. (**A**) Western blot of β-actin, total actin and γ-actin in parental HT1080 (Par) and 3 independent heterozygous FLAG-β-actin HT1080 clones (FLAG/WT). (**B**) Quantification of β-actin protein expressed by the WT allele and the FLAG allele as shown in **A** and normalized to γ-actin. (**C**) Representative western blot showing β-actin expression in parental HT1080 (Par) and homozygous ALFA-β-actin HT1080 cells (ALFA/ALFA). (**D**) Quantification of the β-actin expression from the western blot shown in **C** and normalized to tubulin. (**E**) Representative widefield images from cells that have a CRISPR/Cas9-mediated knock-in of AU1, AU5-t, FLAG or ALFA tag in β-actin. Cells were labeled for F-actin and an antibody/nanobody against the respective tag to visualize F-actin (magenta) and the tagged β-actin (green). Scale bar: 20 µm. Scale bar zoom: 5 µm. (**F**) Colocalization analysis of the microscopy results in E showing the Pearson’s coefficient for each of the internally tagged actins. Individual data points indicate single cells and in total 15 cells from 2 independent experiments were included in the analysis. Bars represent the mean value, and error bars represent standard error of mean (SEM). (**G**) Representative western blot of G-actin and F-actin fraction in heterozygous FLAG-β-actin HT1080 cells that were left untreated or treated with latrunculin A. (**H**) Quantification of the F/G-actin ratio for β-actin expressed by the WT allele and FLAG allele from the western blots shown in **G**. Bars represent the mean value, and error bars represent standard error of mean (SEM).

Next, we assessed the incorporation of the tagged actins into the cytoskeleton by performing immunofluorescence labeling followed by widefield microscopy (**Fig. 2E**). Pearson colocalization analysis demonstrated that all the knock-in tagged actins have a strong overlap with F-actin, indicating that they are well incorporated in actin filaments (**Fig. 2F**). Moreover, high resolution images of actin in lamellipodia and nonmuscle myosin IIA in stress fibers in the homozygous ALFA-β-actin cells indicated that these common actin-based structures had a similar architecture as compared to parental cells (**Fig. S7**). Since the ALFA tag allows intracellular detection of the tagged actins in living cells, we also performed live-cell imaging with the ALFA-β-actin cells. For this, we overexpressed ALFA-Nb-mScarlet in the ALFA-β-actin cells and evaluated its colocalization with F-actin by co-transfecting Lifeact-GFP and determining the Pearson’s coefficient at multiple time points (**Fig. S8, Suppl. Movie 1**). This demonstrated that, also in living cells, there is a very high correlation (R^2^=>0.8) between the fluorescence intensity of the tagged actins and the Lifeact-GFP signal (**Fig. S8B**). This further supports the notion that the T229/A230 internally tagged β-actin is properly assembled into actin filaments.

To corroborate our microscopy results, we sought to biochemically determine the F/G-actin ratio of the tagged and wildtype actin and for this, we used the heterozygous FLAG-β-actin and homozygous ALFA-β-actin cells. The results from these experiments demonstrated that the F/G-actin ratio for FLAG-β-actin and ALFA-β-actin was indistinguishable from wildtype actin (**Fig. 2G-H, Fig. S9**), indicating that the internally tagged β-actin was normally integrated into actin filaments.

Together, these results in fixed and living cells with multiple epitope tags suggest that the T229/A230 residue pair in actin is a versatile position for epitope tagging with only a minor impact on actin expression and no measurable effect on the ability of actin to integrate into filaments.

### Internally tagged actins interact with cofilin, profilin and formins DIAPH1 and FMNL2

To study the molecular interactions of the internally tagged β-actin, we performed a co-immunoprecipitation assay and western blot analysis using FLAG-β-actin and ALFA-β-actin. Since coimmunoprecipitation of actin only allows the investigation of monomeric G-actin interactors, we evaluated the binding of the well-established G-actin binding proteins cofilin and profilin. From the results, we first concluded that both FLAG- and ALFA-β-actin could be immunoprecipitated from the lysates (**Fig. 3A-C**), thereby confirming the availability of the epitope tags under native conditions. More importantly, we could also demonstrate that FLAG-β-actin (**Fig. 3A-B**) as well as ALFA-β-actin (**Fig. 3C**) still associate with cofilin and profilin, indicating that the internally tagged actins maintain their ability to bind to these important actin regulators. For ALFA-β-actin, we further investigated its ability to bind to the actin nucleating formins DIAPH1 and FMNL2, which we also expect to be co-immunoprecipitated with actin under these conditions. The results from these experiments show that both DIAPH1 and FMNL2 associate with ALFA-β-actin (**Fig. 3D**), strongly suggesting that the internal tag does not prevent the binding of actin with formin family members.

**Fig. 3.**
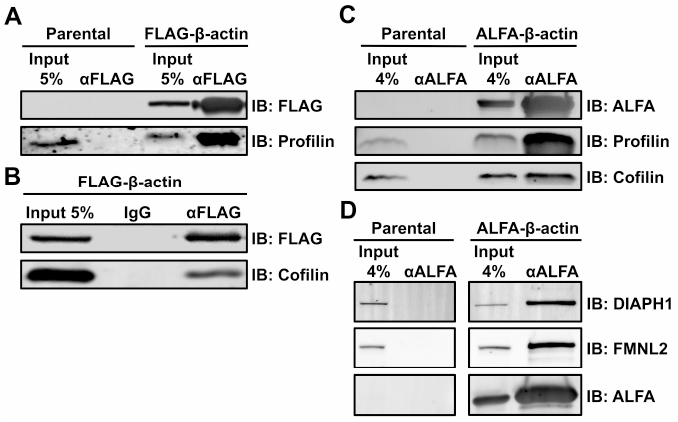
FLAG- and ALFA-β-actin interact with profilin, cofilin and formin family members DIAPH1 and FMNL2. (**A**) Representative western blot showing the co-immunoprecipitation of FLAG-β-actin and profilin using an anti-FLAG antibody in the FLAG-β-actin HT1080 cells. Co-immunoprecipitation performed on parental HT1080 was included as a control. (**B**) Representative western blot showing the coimmunoprecipitation of FLAG-β-actin and cofilin using an anti-FLAG antibody. IgG was included as a negative control. (**C**) Representative western blot showing the co-immunoprecipitation of ALFA-β-actin and profilin and cofilin using an anti-ALFA nanobody in the ALFA-β-actin HT1080 cells. Co-immunoprecipitation performed on parental HT1080 was included as a control. (**D**) Representative western blot showing the co-immunoprecipitation of ALFA-β-actin, mDia1 and FMNL2 using an anti-ALFA nanobody in the ALFA-β-actin HT1080 cells. Co-immunoprecipitation performed on parental HT1080 was included as a control.

### Actin retrograde flow and cell proliferation and migration are not affected by β-actin internal tagging

Fluorescent fusions of actin are known to affect actin retrograde flow at the cell front, likely due to the large fluorescent reporter tag (16). To evaluate whether actin retrograde flow is unperturbed by introducing the internal tag at position T229/A230, we determined the actin flow at lamellipodia using live cell imaging (**Fig. 4A**). For this, we transfected ALFA-β-actin cells with Lifeact-GFP or ALFA-Nb-GFP and performed time lapse imaging with Airyscan super-resolution microscopy. Of note, expression of the ALFA-Nb-GFP did not influence lamellipodia architecture and was similar under all conditions (**Fig. S10**). Parental cells transfected with Lifeact-GFP were included as a control since the expression of Lifeact has been demonstrated to not affect the actin retrograde flow at the cell front (16). Importantly, we observed actin flow at lamellipodia in all of the conditions indicating no gross defects in the formation of these structures by the internal ALFA tag (**Suppl. Movie S2-4**). Moreover, by quantitative analysis of the kymographs from the time-lapse videos, we demonstrate that there are no significant differences in actin flow at lamellipodia between any of the investigated conditions (**Fig. 4B-C**). These results strongly suggest that actin dynamics is not disturbed by the internal tag in β-actin or the overexpression of the ALFA-Nb-GFP.

**Fig. 4.**
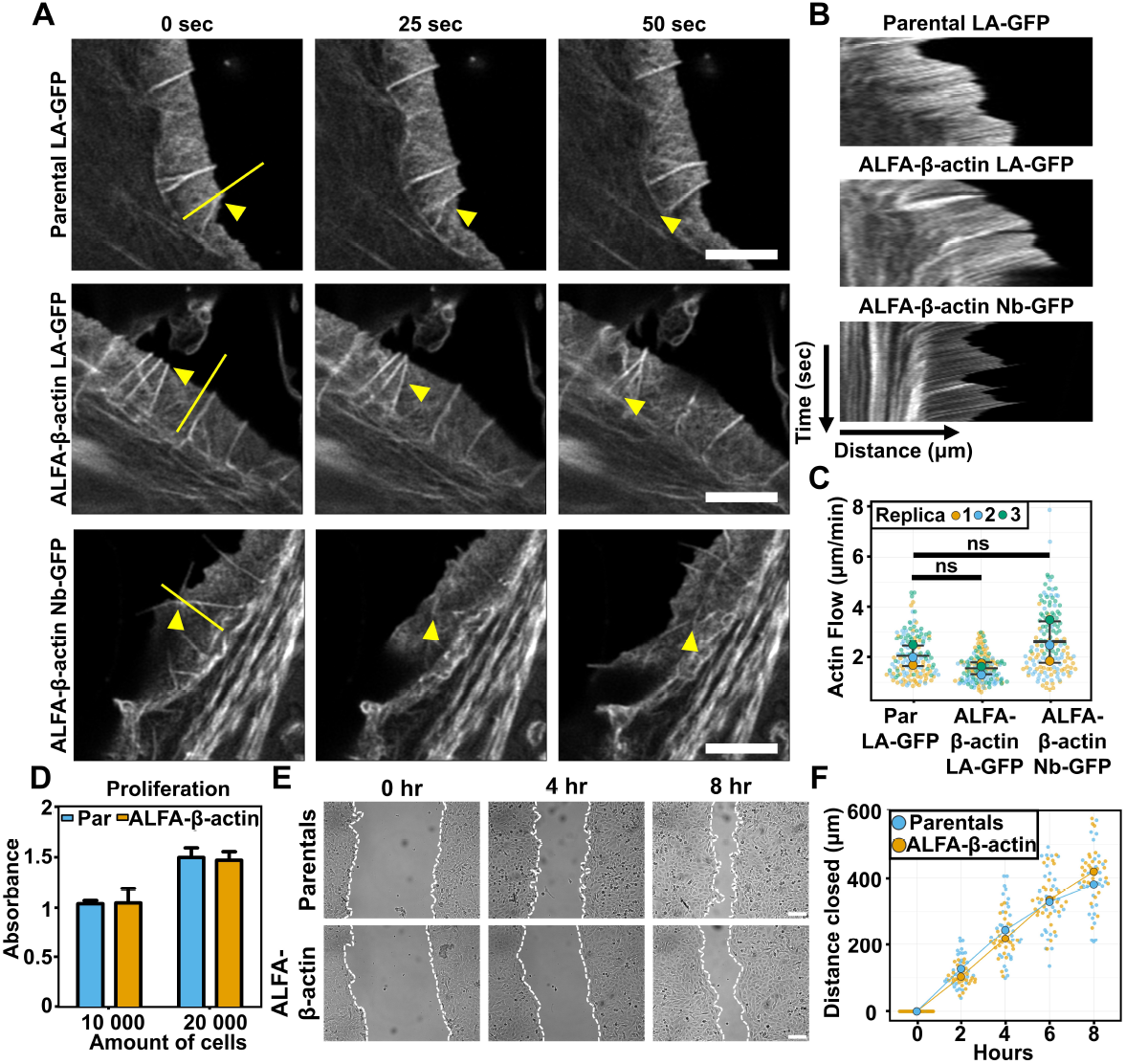
Actin retrograde flow and cell proliferation and migration are not affected by β-actin internal tagging. (**A**) Representative airyscan images of HT1080 parental cells transfected with Lifeact-GFP (LA-GFP), HT1080 ALFA-β-actin cells transfected with Lifeact-GFP and HT1080 ALFA-β-actin cells transfected with ALFA-Nb-GFP. Shown are three stills at timepoint 0, 25 and 50 seconds and the yellow triangles indicate actin features that display retrograde flow. Yellow line indicates the position of kymographs shown in **B**. The full movies are available as Suppl. Movies 2-4. Scale bar: 4 µm. (**B**) Representative kymographs of parental-LA-GFP, ALFA-β-actin-LA-GFP and ALFA-β- actin-Nb-GFP as indicated by the yellow line in **A**. (**C**) Quantification of the actin retrograde flow (µm/min) in parentals-LA-GFP, ALFA-β-actin-LA-GFP, ALFA-β-actin-Nb-GFP. Large datapoints represent the average for each experiment and the small datapoints represent individual kymographs. The bars show the median and the error bars represents standard deviation. Statistical analysis was performed using unpaired Welch’s t-test. Parentals-LA-GFP vs ALFA-β-actin-LA-GFP *P*=0.16. Parentals-LA-GFP vs ALFA-β-actin-Nb-GFP *P*=0.38. (**D**) Quantification of an MTT proliferation assay performed on parental and ALFA-β-actin HT1080 cells. Bars represent the mean value, and error bars represent standard error of mean (SEM) of 2 experiments. (**E**) Representative widefield images of parental and ALFA-β-actin HT1080 cells at time point 0 hr, 4 hr and 8 hr after scratch induction. Scale bar: 30 µm. (**F**) Quantification of the wound healing assay shown in **E** indicating the distance closed in µm over time in parental (Par) and ALFA-β-actin HT1080 cells. Large data points represent the mean of 3 experiments and the small data points represent the quantification of the individual images. 10 images per condition were acquired per experiment.

To demonstrate that the internal tag does not influence cellular processes that are crucially dependent on proper actin function, we evaluated the ability of ALFA-β-actin cells to proliferate and migrate as compared to parental HT1080 cells. To assess cell proliferation, we performed a MTT assay and observed no differences in the the proliferation rate between ALFA-β-actin and parental HT1080 cells (**Fig. 4D**). To assess cell migration, we performed a wound closure assay and demonstrate that the migration rate of the ALFA-β-actin cells is comparable to the parental cells, suggesting there are no major defects in migration speed due to the internal ALFA tag (**Fig. 4E-F**).

Together, these results indicate that actin dynamics as well as major actin-dependent cellular functions are largely unaffected when actin is tagged at postion T229/A230.

### Tagged actin variants recapitulate differential isoform distribution in mammalian cells and yeast

So far, our results strongly suggest that the T229/A230 position in actin is a permissive position for non-disruptive epitope tag integration. We term this internal tagging strategy “IntAct” and propose that it can be used to study the molecular principles of actin isoform specificity in biological processes across species. To demonstrate that the tagged actins recapitulate the behaviour of wildtype isoforms, we first sought to investigate the distribution of cytosolic isoforms in HT1080 cells. For this, we used both the ALFA-β-actin and ALFA-γ-actin knock-in cells that we had generated using CRISPR/Cas9. Since it has been shown before that β- and γ-actin display a differential cellular distribution (11, 38), we evaluated whether the localization of these isoforms is similar in parental and IntAct HT1080 cells. For this, we labelled parental cells with isoform-specific antibodies, and the IntAct cells with an anti-ALFA nanobody as well as phalloidin for normalization. Cells were imaged with Airyscan microscopy and the distribution of actin isoforms was evaluated through ratio calculations. In line with previous observations, we first found that in the parental HT1080 cells, β-actin is enriched at the cell cortex and γ-actin is enriched in stress fibers (**Fig. 5A-C, Fig. S11A**). More importantly, we clearly noted that also in the IntAct HT1080 cells, β- and γ-actin displayed a differential distribution with β-actin being more strongly present at the cortex and γ-actin at the stress fibers (**Fig. 5B-C, Fig. S11B**). Together, these results indicate that isoform-specific properties remain preserved after internal tagging at position T229/A230.

**Fig. 5.**
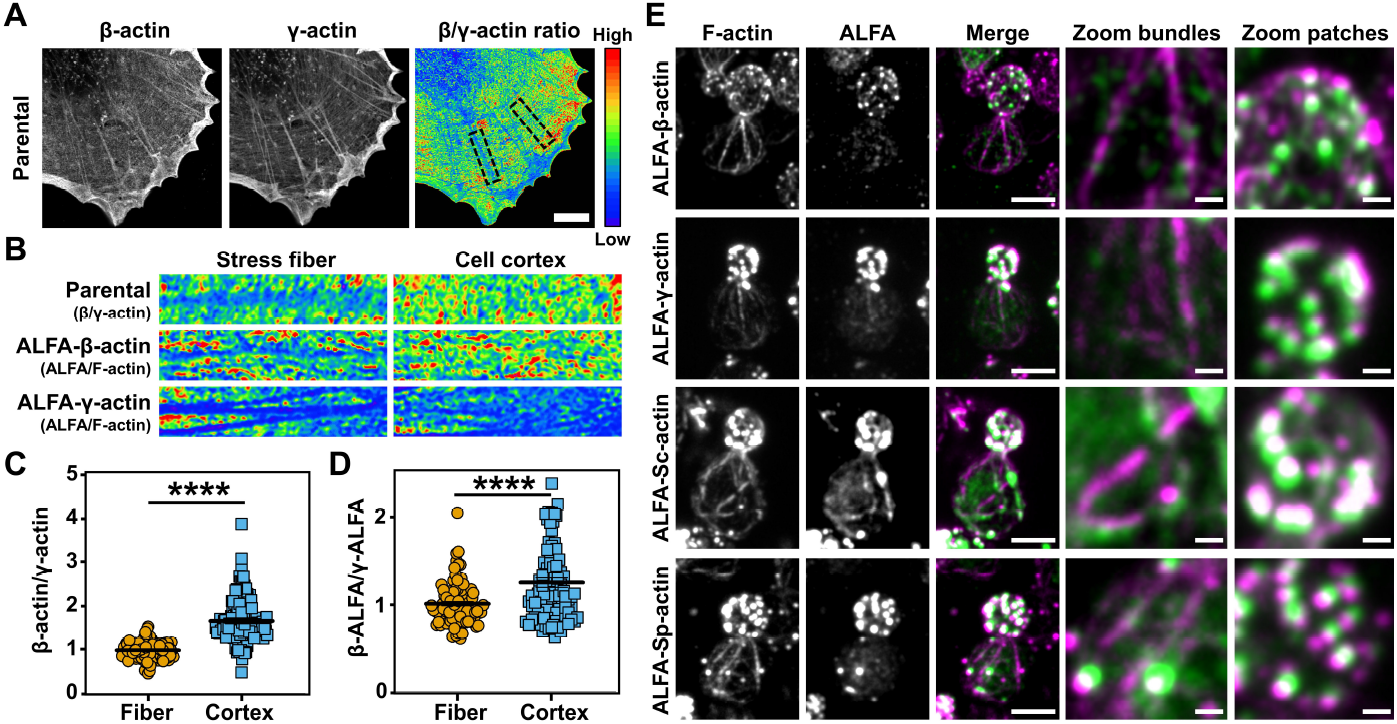
Tagged actin variants recapitulate differential distribution in mammalian cells and yeast. (**A**) Representative Airyscan images of parental HT1080 stained for β-actin (green) and γ-actin (magenta) and a 32-Color ratio image of β-actin divided by γ-actin. Scale bar: 5 µm. (**B**) Representative ratio images of selected regions of interest of stress fibers (left column) and cell cortex (right column) in HT1080 parentals, β- and γ-actin IntAct cells. HT1080 parental cells ratio image was made by dividing the β-actin channel over the γ-actin channel. β- and γ-actin IntAct cell ratio image were made by dividing the ALFA tag channel over the F-actin channel. (**C**) Quantification of the ratio images of HT1080 parentals in **B**. Ratios were normalized against the stress fibers. Per cell, 10 datapoints were collected and in total 15 cells were analyzed over 2 independent experiments. Middle line represents the mean value, and error bars represent standard error of mean (SEM). Statistical analysis was performed using an unpaired t-test *P*=<0.0001. (**D**) Quantification of the ratio images of IntAct cells in **B**. Ratios were first normalized against the F-actin and then de ratio ALFA-β-actin was divided by the ratio of ALFA-γ-actin. Per cell, 10 data points were collected and in total 10 cells were analyzed over 2 independent experiments. Middle line represents the mean value, and error bars represent SEM. Statistical analysis was performed using an unpaired t-test *P*=<0.0001. (**E**) Representative montages of yeast cells expressing β-IntAct, γ-IntAct, Sc-IntAct and Sp-IntAct. Cells were fixed and stained for F-actin (magenta) and ALFA tag nanobody (green). Scale bar: 3 µm, Scale bar zoom: 0.5 µm.

Next, we wondered if our labeling strategy can be applied to label actin in other species. To address this question, we constitutively expressed IntAct ALFA-actins from yeast (*Saccharomyces cerevisiae* and *Schizosaccharomyces pombe*) and human (β- and γ-actin) as an additional copy in a *S. cerevisiae* strain that constitutively expresses a mNeonGreen-ALFA nanobody fusion protein (ALFA-Nb-mNG) (39) (**Fig. S12A-B**). Spot assays of yeast co-expressing IntAct actins and ALFA nanobody showed similar growth rates as the wild type (**Fig. S12C**). Live cell imaging revealed that all IntAct actins displayed dynamic foci-like localization abundant in the growing bud and moved to the bud neck before cytokinesis (**Fig S13A**). This localization pattern resembles that of cortical actin patches in *S. cerevisiae* (40, 41), something which we confirmed by a costaining for F-actin (**Fig. S13B**). These results establish that IntAct actins incorporate into cortical actin patches which consist of branched actin filaments nucleated by the Arp2/3 complex. Consistent with this conclusion, CK666 treatment abolished the actin patch localization of IntAct actins (**Fig. S13C**).

Notably, we did not observe mNG signal into unbranched actin cables which are elongated by formins (Bnr1 and Bni1) into the bud and mother cytoplasm. Since the incorporation of tagged actins into both branched and unbrached actin filaments appeared unperturbed in mammalian cells, we hypothesized that this could be a result of steric hindrance between the ALFA-Nb-mNG and the yeast formin proteins rather than a result of the internal tag itself. To test this, we expressed IntAct ALFA-actins in a wild type *S. cerevisiae* strain and assessed their localization with the actin cytoskeleton. Importantly, although the level of incorporation into actin cables varied among the actin variants, we observed incorporation of IntAct actins into both cortical actin patches and cytoplasmic actin cables in the bud (nucleated by formin Bni1) and mother (nucleated by formins Bnr1 and Bni1) cytoplasm (**Fig. 5E, Fig. S14**). Specifically, *S. cerevisiae* ALFA-actin displayed the greatest degree of cable signal as expected due to species compatibility, while *S. pombe* ALFA-actin showed no detectable localization to actin cables. Furthermore, human ALFA-γ-actin showed weak but discernible localization to actin cables while human ALFA-β-actin only showed localization to cortical actin patches. Together, these results suggest that it is rather the association of ALFA-Nb-mNG with the ALFA-actin monomers than the tag itself that is responsible for the lack of incorporation of tagged actins into actin cables. Furthermore, we conclude that internally tagging actin at the T229/A230 position results in a functional actin protein with the ability to incorporate into linear and branched actin filaments in both human and yeast cells, indicating that our IntAct internal tagging strategy can be used to study actin diversity and functionality across species.

## Discussion

In this manuscript, we present a strategy to internally tag actin to study the molecular interactions and dynamics of actin isoforms. At this point, we can only speculate as to why the T229/A230 position seems permissive for manipulation. The T229/A230 residue pair is located in subdomain 4 and is part of a region that has been termed the V-stretch due to the high structural variation that this region exhibits in molecular dynamics simulations of F-actin (42). To the best of our knowledge, the V-stretch domain has no explicitly described interactions with actin-binding proteins (ABPs), either to G-actin monomers or to F-actin filaments. Specifically, the majority of ABPs that associate with monomeric actin including profilin (43) and gelsolin (44) bind to the barbed end of the actin molecule on subdomains 1 and 3 (45), which is relatively distant from the T229/A230 position. On the other hand, many F-actin associated molecules including myosin (46) and cofilin (47) bind to the so-called outer domain of the actin molecule which comprises part of subdomain 1 and 2 which is also not adjacent to the V-stretch in subdomain 4. Finally, unlike other variable regions such as the D-loop in subdomain 2, the V-stretch is not involved in interactions between monomers in actin filaments (48). Interestingly, an alanine mutagenesis scan of the entire β-actin protein further demonstrated that the V-stretch has a high structural plasticity since the alanine mutants covering this region were not impaired in their folding capacity or their binding to the actin-binding proteins DNAse I, adseverin, Thymosin β4 and CAP (49). It must be noted though that the T229/A230 residue pair is very close to the proposed binding interface of the actin-binding protein nebulin, and therefore possibly also the related protein nebulette (50). Nonetheless, since nebulin and nebulette are exclusively present in skeletal and cardiac muscle, respectively, we expect this will not be an obstacle for studying the non-muscle and smooth muscle actin isoforms. Future investigations using internally tagged α-skeletal or α-cardiac actin, however, need to carefully control for the possibility that the binding with nebulin or nebulette is influenced by the use of the T229/A230 residue pair.

A previous attempt to internally tag actin with a tetra-cysteine tag in yeast demonstrated that, out of eight different internal sites, only the S232/S233 position allowed weak assembly of actin into filament cables (31). We demonstrate here that IntAct actins are integrated into yeast filament cables, although the extent of integration depends on the species and isoform variant that is used. This differential incorporation into actin structures shown by different IntAct proteins is likely an outcome of hindered interaction with key ABPs arising due to evolutionary divergence from the actin protein sequence of *S. cerevisiae*. Similar isoform and species dependent differences have been demonstrated in a recent study where various actin proteins showed differential recruitment to linear and branched actin structures upon expression in *S. cerevisiae* (51). It would be interesting to use IntAct actins to dissect the differential binding partners and molecular mechanisms that explain this difference. The fact that the internal position identified by us is extremely close to the S232/S233 residue pair that was previously selected further indicates that this particular region in the actin molecule is relatively permissive for manipulation. The fact that the internal position identified by us is extremely close to this S232/S233 residue pair further indicates that this region is relatively permissive for manipulation. Interestingly, in our screen, we also included the S232/S233 residue pair as well as the other two adjacent positions, i.e. A230/A231 and A231/S232. These internally tagged variants, however, were not as well assembled into filaments as the T229/A230 variant, suggesting very specific structural requirements for actin internal tagging.

We show that in cells from human and yeast, IntAct actins are integrated in F-actin based structures that are composed of branched filaments such as lamellipodia and actin patches as well as unbranched filaments such as filopodia, stress fibers and actin cables. For lamellipodia, we also demonstrate that rearward treadmilling, which is indicative for polymerization speed, is not different between cells that express wildtype or IntAct β-actin. It is well established that the assembly of branched actin filaments is controlled by Arp2/3 complex formins, while the assembly of unbranched filaments is controlled by formins (52). We therefore conclude that internal tagging of actin at position T229/A230 does not prevent the binding to these different classes of actin nucleating and elongation factors, something which is supported by the binding of formins to IntAct β-actin. Although our findings strongly suggest that the nucleation and polymerization of actin is largely unaffected, future in vitro reconstitution experiments will be required to precisely dissect to what extent the biochemical properties of IntAct actins are similar to those of wildtype actins. For this, recent developments in actin production and purification systems such as Pick-ya actin could be very instrumental (53, 54).

An important difference between our results in yeast and human cells is the ability to visualize the ALFA-tagged actins in living cells. In human cells, the nanobody allowed visualization of various types of actin-based structures including filopodia, lamellipodia and stress fibers while in yeast, only the branched actin patches were visible and the visualization of unbranched actin cables was hampered. It remains to be determined whether this discrepancy is due to improper incorporation of ALFA-actin in the presence of the nanobody or because of steric hindrance of the Nb-ALFA-mNG fusion protein to localize to actin cables. Our finding that ALFA-tagged actin has the ability to be integrated in unbranched actin cables in the absence of the nanobody at least indicates that the tag itself is likely not the reason for our inability to visualize actin cables in living cells. For potential future live cell imaging experiments to study yeast actin cables, optimization of other small tag/fluorophore combinations is therefore required. It should be further noted that in this study, we have only established one small tag/fluorophore combination for live cell targeting of the tag in living cells, namely the ALFA tag. While our findings extend the applicability of the ALFA-tag system to visualize homologous proteins with known tagging issues, something which for example could also overcome labeling strategies for tubulin isoforms, for future experiments that require simultaneous targeting of two, or more, isoactins in the same cell, additional tag/fluorophore combinations should be optimized. For this, we expect that novel bottom-up design approaches for nanobody-based tags, similar to the ALFA tag design (35), is necessary.

In conclusion, our results demonstrate that the T229/A230 residue pair allows internal tagging of actin, which can be used to study the localization and dynamics of specific actin isoforms. Also, we show that this internal tagging strategy enables the investigation of molecular interactions of monomeric actin using standard pulldown assays. Interestingly, since our approach involves intracellular targeting of specific actin variants, we envision that fusing the anti-ALFA nanobody with peroxidases (e.g. APEX2 (55)) or biotin ligases (e.g. TurboID (56)) will allow the future investigation of isoactin-specific molecular interactions of both monomeric and filamentous actin. Furthermore, we also demonstrate the possibility of tagging actin at position T229/A230 in yeast actins, suggesting that our IntAct approach can be applied across species to study differences in isoactins. Finally, we envision that, by tagging mutant actin variants, our approach could also open avenues to unravel the disease causing mechanisms of a wide variety of actinopathies (4), for which currently no strategy is available.

### Experimental Procedures

#### Cell Culture

HT1080 fibrosarcoma cells were used for the overexpression of internally tagged actins and to generate the internally tagged cell lines. Cells were cultured in 1x DMEM + 4.5 g/L D-Glucose, NEAA (Gibco, Lot#2246377) and supplemented with 10% (vol/vol) fetal bovine serum (FBS), 1X Glutamax (Gibco, 2063631), 1 mM Sodium Pyruvate (Gibco, 2010382) and 0.5X Antibiotic-Antimycotic (Gibco, 15240-062). RPE1 cells were used for the overexpression of internally tagged actins. RPE1 cells were cultered in advanced DMEM/F-12 + non-essential amino acids + 110mg/L Sodium Puruvate (Gibco, Lot#12634010) supplemented with 10% (vol/vol) fetal bovine serum (FBS) and 1X Glutamax (Gibco, 2063631). All cell lines were cultured and kept at 37°C with 5% CO2.

#### Yeast strains

All yeast strains used in this study are listed in Supplementary Materials (**Suppl. Table 1**). Yeast strains were constructed using protocols previously described (57).

#### Antibodies and reagents

The following primary antibodies were used: anti-β-actin (#MCA5775GA, Bio-Rad Laboratories, 1:100 for immunofluorescence, 1:1000 for western blot), anti-γ-actin (#MCA5776GA, Bio-Rad Laboratories, 1:100 for immunofluorescence, 1:1000 for western blot), anti-Flag (#F1804-1MG, Sigma Aldrich, 1:1000 for western blot), anti-AU1 (#NB600-452, Novus biologicals, 1:100 for immunofluorescence), anti-AU5 (#NB600-461, Novus biologicals, 1:100 for immunofluorescence), anti-profilin (#3246, Cell Signaling, 1:1000 for western blot), anti-cofilin (#5175P, Cell Signaling, 1:1000 for western blot), anti-DIAPH1 (#610848, BD Science, 1:1000 for western blot), anti-FMNL2 (#HPA005464, Sigma Aldrich, 1:1000 for western blot), anti-Myosin IIA Heavy chain (#909801, Biolegend, 1:100 for immunofluorescence), antitubulin (homemade antibody, 1:4000 for western blot), anti-aSMA (#19245S, Cell signaling, 1:1000 for western blot). Actin was stained with anti-actin (#A2066, Sigma Aldrich, 1:4000 for western blot), Alexa-488-conjugated phalloidin or Alexa-568-conjugated phalloidin (Life Technologies, 1:200), ALFA tag was stained with HRP-conjugated sdAb anti-ALFA primary antibody (#N1505, NanoTag biotechnologies, 1:1000 for western blot), anti-ALFA-atto488 conjugate or anti-ALFA-Alexa647 conjugate (#N1502 NanoTag biotechnologies, 1:100 for immunofluorescence). Secondary antibodies conjugated to Alexa647, Alexa568, or Alexa 488 were used (Life Technologies, 1:400 for immunofluorescence).

#### Generation of overexpression constructs

All overexpression constructs with the internally tagged actins were generated by Gibson assembly (New England Biolabs). Briefly, two PCRs were performed per construct to generate a DNA fragment upstream of the tag and a fragment downstream of the tag. Primers used for the PCR reactions are given in the Supplementary Materials (**Suppl. Table 2**). Both fragments contained the DNA coding for the tag which functions as an overlapping sequence in the Gibson assembly reaction. The pcDNA3.1 vector backbone was linearized using restriction enzymes HindIII and NheI and 100 ng of vector was used in every Gibson assembly. PCR fragments were added in a 1:6 vector:insert molar ratio and all assembly reactions were incubated for 50 degrees for 50 min. Half the product was transformed into Top10 competent bacteria and clones were screened for the correct vectors.

#### Generation of knock-in cell lines

gRNAs and HDR templates used for the generation of FLAG-, AU1-, AU5-t- and ALFA-knock-in cells are given in the Supplementary Materials (**Suppl. Table 2**). Lipofectamine 2000 (ThermoFisher, ref. 11668027) was used to transfect the HT1080 cells. To increase the efficacy of the knockin approach we applied a coselection procedure using ouabain as described previously (36). The gRNA and HDR template for mutating the ATP1A1, which leads to ouabain resistance, are given in the Supplementary Materials (**Suppl. Table 2**). Flow cytometery for the respective tags was performed two weeks after the initial transfection to determine the number of positive cells. Subsequently, treatment with 0.75 µM ouabain was started and after two weeks, single clones were generated from the selected cells. Positive clones were selected based on intracellular FACS staining and were further used for immunofluorescence and western blot.

#### Generation of yeast constructs

All plasmids used in this study are listed in Supplementary Materials (**Suppl. Table 4**). The IntAct DNA sequence for *Saccharomyces cerevisiae* (Sc-IntAct) and *Schizosaccharomyces pombe* (Sp-IntAct) were synthesized by GeneArt (Life Technologies). Sequences for human β-IntAct and human γ-IntAct were amplified from plasmids used for overexpression. All plasmids were constructed using NEBuilder Hifi (New England Bio-Labs).

#### Immunofluorescence mammalian cells

All steps were performed at room temperature. Cells were seeded on coverslips and fixed using 4% PFA for 10 min. Permeabilization was performed with 0.1% Triton X-100 for 5 min. After washing with 1x PBS, the samples were blocked with 20 mM PBS+glycin. Primary Ab incubation was performed for 1 hr. Subsequently, samples were washed 3x with 1x PBS and incubated with the appropriate secondary Ab for 1 hr in the dark. The samples were then washed 2x with 1x PBS and 1x with MilliQ. After these washing steps the samples were sealed in Mowiol and dried overnight.

#### Immunofluorescence yeast

Immunofluorescence was performed using a modified version of a protocol described previously (58). Briefly, yeast strains were grown overnight at 25°C in YPD broth. The overnight culture was diluted and allowed to grow until mid-log phase. Cells were fixed with 4% formaldehyde for 60 minutes at 25°C, washed twice with 1x PBS, and finally resuspended in 200µL of 1.2M Sorbitol Phosphate-Citrate (SPC) buffer (1.2M Sorbitol, 1M K2HPO4, 1M Citric acid). 25µL of Long-Life Zymolase (GBiosciences) was added to digest the yeast cell wall and the suspension was incubated with mild shaking at 37°C for 60 minutes. The cells were then washed twice with ice-cold SPC buffer and incubated with 500µL of blocking buffer 2% Bovine Serum Albumin (BSA) + 0.1% Triton X-100 in PBS) at room temperature for 15 minutes with shaking. The cells were pelleted and resuspended in 500µL of Antibody Dilution Buffer (1% BSA + 0.05% Triton X-100 in PBS) containing FluoTag-X2 anti-ALFA-Alexa647 at a final dilution of 1:500. The cell suspension was then incubated overnight with rotation at 4°C. Next day, cells were washed twice with 1xPBS and finally resuspended in 20µL of 1xPBS. 5µL of the final cell suspension was mounted on a glass bottom dish coated by 6% Concanavalin A. Phalloidin staining of yeast actin structures was done as per previously described protocols (59, 60). Briefly, cells were grown at 25°C till mid-log phase and fixed with 4% paraformaldehyde. The cells were washed thrice with 1xPBS and labelled phalloidin was added to a final concentration of 0.4µM (in 50µL of 1x PBS) and the tubes were kept in a rotating shaker overnight at 4°C. The cells were washed twice with 1x PBS on the next day and seeded on a concanavalin A coated glass-bottom dish. Images were acquired using Andor Dragonfly 502 spinning-disk confocal setup consisting of a Leica Dmi8 fully motorized inverted microscope equipped with Andor Sona scMOS camera or Olympus FV3000 point-scanning confocal setup consisting of an Olympus IX83 fully motorized inverted microscope equipped with high-sensitivity GaAsP photomultiplier tube (PMT) detectors.

#### Imaging

Imaging was performed on a Zeiss LSM900 laser scanning confocal microscope and a Leica DMI6000 epifluorescence microscope. Images on the LSM900 were acquired using a 63x 1.4 NA oil objective. Alexa488 was excited at 488 nm and emission light was detected between 490-575 nm. Alexa568 was excited at 561 nm and emission was detected between 555-700 nm. Alexa647 was excited at 640 nm and emmission was detected between 622-700 nm. Raw images were processed using the Zeiss Zen 3.1 blue edition software. Images on the Leica DMI6000 were acquired with an HC PL APO 63x 1.40NA oil objective and a metal halide lamp. Alexa488 was excited between 450-490 nm and emmision light was detected between 500-550nm. Alexa568 was excited between 540 nm and emmision light was detected between 567-643 nm.

#### Live-cell imaging mammalian cells

Parental and/or ALFA β-actin HT1080 cells were seeded in WillCo wells (WillCo Wells B.V.). The next day, cells were transfected with Lifeact-GFP, Nb-ALFA-GFP or Nb-ALFA-mScarlet together with Lifeact using Lipofectamine 2000 (Invitrogen, lot#1854327). Prior to imaging, DMEM was replaced by imaging medium (HBSS, Ca/Mg, 5% FCS, 25 mM HEPES) and incubated for approximately 10 min. Live cell imaging was performed using a Zeiss LSM880 with Airyscan and data was acquired using a 63x 1.4 NA oil objective. During single color live-cell imaging, Lifeact-GFP and Nb-GFP was excited using a mass beam splitter (MBS) 488 and emission light was collected using a 495-550 band pass/570 long pass (LP) filter. Sequentially imaging was done for Lifeact-GFP together with Nb-mScarlet. Lifeact-GFP and Nb-mScarlet were excited using an MBS 488/561 and emission light was collected using a BP 420–480/BP 495–550 for Lifeact-GFP and emission light was collected using a BP 570-620 + LP 645 for Nb-mScarlet. Time series were collected with a frame interval of 5 s for actin treadmilling and 15 s for colocalization of Lifeact with Nb-mScarlet. Raw images were processed using the Zeiss Zen 2.1 Sp1 software. Image series were analyzed using ImageJ and the Pearson correlation coefficient was calculated using the ImageJ tool Coloc2 (PSF: 3, Costes randomizations 10).

#### Live-cell imaging of yeast cells

Yeast cells were grown at 25°C to mid-log phase in Synthetic Complete (SC) medium and plated on a glass-bottom dish coated by 6% Concanavalin A. For CK666 treatment, DMSO (solvent control) or CK666 was added to a final concentration of 200µM in the media contained in the glass bottom dish and the prepared dish was taken for live cell imaging. Images were either acquired using Andor Dragonfly 502 spinning-disk confocal setup consisting of a Leica Dmi8 fully motorized inverted microscope equipped with Andor Sona scMOS camera or Olympus FV3000 point-scanning confocal setup consisting of an Olympus IX83 fully motorized inverted microscope equipped with high-sensitivity GaAsP photomultiplier tube (PMT) detectors.

#### Western blot

For western blot, cells were lysed with 2x Laemmli (5 ml 0.5 M Tris pH 6.8, 8 ml 10% SDS, 4 ml Glycerol, few grains bromophenol blue, 2 ml 2-mercaptoethanol and 1 ml MilliQ). Samples were loaded onto 10% or 15% SDS-PAGE gels for separation. Separation was accomplished by running for approximately 2 hrs at 100 V in 1x Running buffer (100 ml 10x TBS, 10ml 10% SDS and 900 ml MilliQ). Proteins were transferred to PVDF membranes for approximately 1 hr at 100 V in transfer buffer (100 ml 10x TBS, 200 ml MeOH and 700 ml MilliQ). Membranes were blocked in 5% milk (ELK milk powder, Fries-land Campina) in TBST (1x TBS, 0.1% Tween20) and incubated with primary antibodies overnight at 4°C while rotating. After three times washing with 1x TBST, membranes were incubated in the dark for 1 hr with secondary antibodies while rotating. Washing with 1x TBST was repeated and subsequently, the protein bands were visualized using a Typhoon FLA 7000 (GE Healthcare). ImageJ was used to analyze the protein bands. For yeast cell lysates, TCA precipitation method was used as described (39), protein was probed with HRP-conjugated sdAb anti-ALFA primary antibody (#N1505, NanoTag biotechnologies).

#### F-/G-actin ratio

Cells were seeded and the next day washed in ice-cold PBS and lysed on ice for 10 minutes with F-actin stabilization buffer (0.1 M PIPES pH 6.9, 30% glycerol, 5% DMSO, 1 mM MgSO4, 1 mM EGTA, 1% Triton X-100, 1 mM ATP, protease inhibitor cocktail (Sigma, 11697498001)). The cells were harvested and spun down for 10 minutes at 1,000 g and 4°C. The supernatant was collected and spun down at 16,000 g for 75 minutes at 4°C to separate the G- and F-actin fractions. The supernatant, containing G-actin, was collected and the pellet, containing F-actin, was solubilized in depolymerization buffer (0.1 M PIPES pH 6.9, 1 mM MgSO4, 10 mM CaCl2, 5 µM cytochalasin D, 1% SDS) or 2x Laemmli (5 ml 0.5 M Tris pH 6.8, 8 ml 10% SDS, 4 ml Glycerol, few grains bromophenol blue, 2 ml 2-mercaptoethanol and 1 ml MilliQ). For the negative control, cells were treated with 1 µM Latrunculin A 30 minutes before lysis to disrupt the F-actin fraction. The F-/G-ratio was determined by western blot analysis.

#### ALFA tag co-immunoprecipitation

Cells were seeded and the following day, the cells were washed five times with ice-cold PBS. Cells were lysed with ice-cold lysis buffer (10 mM Tris, 150 mM NaCl, 2 mM MgCl2, 2 mM CaCl2, 1% Brij-97), supplemented with 1x protease inhibitor cocktail (Roche, 11697498001) and 0.1 mM PMSF (Sigma-Aldrich, P7626-5G). Cell lysates were centrifugation at 16,000 g for 60 minutes at 4°C. For ALFA tag pulldowns, ALFA-Selector ST beads (NanoTag biotechnologies, N1510) were washed twice with lysis buffer. For input 4% or 5% of the clarified lysate was collected as positive control and diluted in 2x Laemmli buffer. The rest of the sample was combined with the beads and incubated for 1 hour at 4°C with rotation. After enrichment, the beads were pelleted by centrifugation for 1 minute at 1,000 g, and the supernatant was collected as negative control. The beads were washed five times with lysis buffer and incubated for 20 minutes with 2x Laemmli buffer supplemented 0.2 µM elution peptide (NanoTag biotechnologies, N1520-L) at RT with subtle shaking. The samples were centrifuged for 1 minute at 3,000 g and the supernatant collected as elute sample.

#### FLAG co-immunoprecipitation

The day before starting the co-immunoprecipitation protocol, BSA- and IgG-coated dynabeads for pre-clearing were prepared by mixing dynabeads slurry (40 µl per sample) with 500 µl PBS/3% BSA or 500 µl PBS with 2.5 µg mouse IgG1 (BioLegend, 400102) respectively. Similarly, BSA-coated dynabeads for the IP itself were prepared by mixing 60 µl dynabeads slurry per sample with 500 µl PBS/3% BSA. FLAG knock-in clones were seeded and when the cells reached full confluency, they were washed once with cold PBS and lysed for 15 minutes at 4°C with 1 ml lysis buffer (1% Brij-97, 10 mM Tris-HCl pH 7.5, 150 mM NaCl, 2 mM MgCl2, 2 mM CaCl2, protease inhibitor cocktail (Sigma, 11697498001)). Cell lysates were collected by scraping and spun down at 16,000 g and 4°C for 75 minutes to remove F-actin. In the meantime, the pre-clear beads were washed once with lysis buffer (with or without 1 mM ATP) and resuspended in a total of 40 µl per sample. The supernatant was pre-cleared for 1 hour at 4°C while shaking, with both the BSA- and IgG-coated beads. After preclearing, the beads were removed and 40 µl of the supernatant was collected as input. The rest of the sample was split into two parts, which was supplemented with lysis buffer to a volume of 2 ml. 6 µg of either mouse anti-FLAG (Sigma, F1804) or mouse IgG1 (BioLegend, 400102) was added and samples were incubated for 1 hour at 4°C under rotation. Meanwhile, the IP beads were washed with lysis buffer (with or without 1 mM ATP) and resuspended in a total of 60 µl per sample. The beads were added and the samples were incubated for an additional 2 hours. After washing five times with washing buffer (1% Brij-97, 10 mM Tris-HCl pH 7.5, 150 mM NaCl, 2 mM MgCl2, 2 mM CaCl2, 1 mM PMSF), the beads were eluted in 100 mM glycine pH 3.0 for 5 minutes while rotating. The samples were neutralized by adding 1/10th volume of 1M Tris-HCl pH 8.5 and loaded on 15% SDS-PAGE gels for western blot analysis.

#### Wound closure assay

Cells were seeded to 100% confluency and a scratch was made with a 200 µl tip from the top to the bottom of the well. After scratching, the cells were washed 1x with PBS and fresh media was added to the cells. The same position was imaged with a Leica DMI6000 epifluorescence microscope after 0 hrs, 4 hrs and 8 hrs. The images were analyzed using the Wound healing size tool plugin in ImageJ (61). First, the edges of the wound were manually annotated and the plugin was subsequently used to measure the average wound width. Using the average wound width, the relative distance closed between every timepoint and timepoint 0 was calculated.

#### MTT proliferation assay

Cells (10.000 or 20.000) were seeded in a 96-well plate. After 16 hrs of incubation, media was replaced by media containing the tetrazolium dye MTT (0.45 mg/ml). After 2 hrs of MTT incubation at 37C in the CO2 incubator, 150 µl DMSO was added to the cells and incubated for 10 min on an orbital shaker until the crystals were dissolved. The absorbance at 560 nm was determined on a microplate reader (iMark microplate absorbance reader, Bio-Rad).

#### Image analysis

All image analysis was performed using ImageJ (62). Pearson correlation coefficient was calculated using the ImageJ tool Coloc2 (PSF: 3, Costes randomizations 10). For the ratio analysis, first a background subtraction was performed. For the parental cells, the β-actin channel was divided by the γ-actin channel which resulted in a β-/γ-actin ratio image. For visualization, the range was set between 0 and 2 and a 32-color LUT was assigned to the image. For each cell, 10 regions (0.5 µm x 0.5 µm) were selected for stress fibers or the cell cortex and the average ratio value of these regions was measured. The ratios were subsequently normalized to the value of the stress fibers. For the IntAct cells, first the individual ratios of ALFA-β-actin or ALFA-γ-actin over total actin were calculated by dividing the ALFA tag signal by the phalloidin signal. For visualization, the brightness/contrast was set between 0 and 2 and for each cell, 10 regions (0.5 µm x 0.5 µm) were drawn for stress fibers or the cell cortex and the average ratio value of these regions was measured. To calculate the β-/γ-actin ratio, ratio image of ALFA-β-actin was divided by the ratio image of ALFA-γ-actin. Subsequently, these ratios were normalized to the stress fibers and used for graphical representation. Angles of branched actin were measured in lamellipodia using the Angle tool in ImageJ. Myosin IIA spacing was measured on stress fibers by drawing a line along the stress fiber of 7.5 µm, which was saved as a region of interest (ROI). The ROI was overlaid in the myosin IIA channel and an intensity plot profile was made. Subsequently the positions of each high intensity peak in the plot profile were saved. The distance between every high intensity peak along a stress fiber was calculated and an average of all the distances were taken as the average myosin IIA spacing along the single stress fiber. This procedure was repeated for at least 30 stress fibers per condition.

#### Data visualization

All graphs were designed in GraphPad Prism 9.0 (GraphPad Software, Inc.) except for Fig. 4C and 4F which were generated with SuperPlotsofData (63).

#### Statistics

The type of statistical test, *n* values, and *P* values are all listed in the figure legends or in the figures. All statistical analyses were performed using Graph Pad Prism or Microsoft Excel, and significance was determined using a 95% confidence interval.

#### Data availability

All primary data supporting the conclusions made are available from the authors on request.

## ACKNOWLEDGEMENTS

We are indebted to NanoTag Biotechnologies GmbH, Göttingen, Germany for providing us with the ALFA nanobody expression construct. The authors further thank the Radboudumc Technology Center Microscopy for the use of their facilities. AD acknowledges GATE Fellowship by IISc. This work was financially supported by SERB SRG grant (SRG/2021/001600) and a Department of Biotechnology-Wellcome Trust India Alliance intermediate fellowship (IA/I/21/1/505633) awarded to S.P., intramural funding of the Radboudumc and by an NWO KLEIN grant (OCENW.KLEIN.494) awarded to K.D..

## AUTHOR CONTRIBUTIONS

E.S. generated the overexpression constructs. S.W., W.B. and W.S. generated the HT1080 knock-in cells. W.B and M.C.Z. performed the coimmunoprecipitation experiments. B.J. performed microscopy experiments. M.C.Z. performed the overexpression experiments, live-cell imaging, proliferation, and migration experiments. A.D. generated the yeast expression constructs and performed yeast live-cell imaging, immunofluorescence, immunoblotting. K.D. and M.C.Z. carried out data analysis. S.P conceived and supervised the yeast experiments. K.D. conceived and supervised the experiments in human cells. K.D., S.P., M.C.Z. and A.D. wrote the manuscript.

## Bibliography

1. L. R. Andrade. Evidence for changes in beta- and gamma-actin proportions during inner ear hair cell life. Cytoskeleton (Hoboken), 72(6):282–91, 2015.

2. K. M. McHugh, K. Crawford, and J. L. Lessard. A comprehensive analysis of the developmental and tissue-specific expression of the isoactin multigene family in the rat. Dev Biol, 148(2):442–58, 1991.

3. D. Tondeleir, D. Vandamme, J. Vandekerckhove, C. Ampe, and A. Lambrechts. Actin isoform expression patterns during mammalian development and in pathology: insights from mouse models. Cell Motil Cytoskeleton, 66(10):798–815, 2009.

4. F. Parker, T. G. Baboolal, and M. Peckham. Actin mutations and their role in disease. Int J Mol Sci, 21(9), 2020. ISSN 1422-0067 (Electronic) 1422-0067 (Linking). doi: 10.3390/ijms21093371.

5. B. J. Perrin and J. M. Ervasti. The actin gene family: function follows isoform. Cytoskeleton (Hoboken), 67(10):630–4, 2010.

6. S. Baranwal, N. G. Naydenov, G. Harris, V. Dugina, K. G. Morgan, C. Chaponnier, and A. I. Ivanov. Nonredundant roles of cytoplasmic beta- and gamma-actin isoforms in regulation of epithelial apical junctions. Mol Biol Cell, 23(18):3542–53, 2012.

7. M. Moradi, R. Sivadasan, L. Saal, P. Luningschror, B. Dombert, R. J. Rathod, D. C. Dieterich Blum, and M. Sendtner. Differential roles of alpha-, beta-, and gamma-actin in axon growth and collateral branch formation in motoneurons. J Cell Biol, 216(3):793–814, 2017.

8. A. Chen, P. D. Arora, C. A. McCulloch, and A. Wilde. Cytokinesis requires localized betaactin filament production by an actin isoform specific nucleator. Nat Commun, 8(1):1530, 2017.

9. A. Chen, L. Ulloa Severino, T. C. Panagiotou, T. F. Moraes, D. A. Yuen, B. D. Lavoie, and Wilde. Inhibition of polar actin assembly by astral microtubules is required for cytokinesis. Nat Commun, 12(1):2409, 2021.

10. V. Dugina, I. Zwaenepoel, G. Gabbiani, S. Clement, and C. Chaponnier. Beta and gamma-cytoplasmic actins display distinct distribution and functional diversity. J Cell Sci, 122(Pt 16):2980–8, 2009.

11. E. Pasquier, M. P. Tuset, S. Sinnappan, M. Carnell, A. Macmillan, and M. Kavallaris. gamma-actin plays a key role in endothelial cell motility and neovessel maintenance. Vasc Cell, 7: 2, 2015.

12. M. Karakozova, M. Kozak, C. C. Wong, A. O. Bailey, 3rd Yates, J. R., A. Mogilner, H. Zebroski, and A. Kashina. Arginylation of beta-actin regulates actin cytoskeleton and cell motility. Science, 313(5784):192–6, 2006.

13. F. Zhang, S. Saha, S. A. Shabalina, and A. Kashina. Differential arginylation of actin isoforms is regulated by coding sequence-dependent degradation. Science, 329(5998):1534–7, 2010.

14. P. Vedula, S. Kurosaka, B. MacTaggart, Q. Ni, G. Papoian, Y. Jiang, D. W. Dong, and Kashina. Different translation dynamics of beta- and gamma-actin regulates cell migration. Elife, 10, 2021.

15. J. A. Cooper. Effects of cytochalasin and phalloidin on actin. J Cell Biol, 105(4):1473–8, 1987.

16. J. Riedl, A. H. Crevenna, K. Kessenbrock, J. H. Yu, D. Neukirchen, M. Bista, F. Bradke, D. Jenne, T. A. Holak, Z. Werb, M. Sixt, and R. Wedlich-Soldner. Lifeact: a versatile marker to visualize f-actin. Nat Methods, 5(7):605–7, 2008.

17. M. J. Schell, C. Erneux, and R. F. Irvine. Inositol 1,4,5-trisphosphate 3-kinase a associates with f-actin and dendritic spines via its n terminus. J Biol Chem, 276(40):37537–46, 2001.

18. B. M. Burkel, G. von Dassow, and W. M. Bement. Versatile fluorescent probes for actin filaments based on the actin-binding domain of utrophin. Cell Motil Cytoskeleton, 64(11): 822–32, 2007.

19. M. Melak, M. Plessner, and R. Grosse. Actin visualization at a glance. J Cell Sci, 130(3): 525–530, 2017. ISSN 1477-9137 (Electronic) 0021-9533 (Linking). doi: 10.1242/jcs.189068.

20. A. Rocchetti, C. Hawes, and V. Kriechbaumer. Fluorescent labelling of the actin cytoskeleton in plants using a cameloid antibody. Plant Methods, 10:12, 2014. ISSN 1746-4811 (Print) 1746-4811 (Linking). doi: 10.1186/1746-4811-10-12.

21. C. R. Schiavon, T. Zhang, B. Zhao, A. S. Moore, P. Wales, L. R. Andrade, M. Wu, T. C. Sung, Y. Dayn, J. W. Feng, O. A. Quintero, G. S. Shadel, R. Grosse, and U. Manor. Actin chromobody imaging reveals sub-organellar actin dynamics. Nat Methods, 17(9):917–921, 2020. ISSN 1548-7105 (Electronic) 1548-7091 (Linking). doi: 10.1038/s41592-020-0926-5.

22. V. Brault, U. Sauder, M. C. Reedy, U. Aebi, and C. A. Schoenenberger. Differential epitope tagging of actin in transformed drosophila produces distinct effects on myofibril assembly and function of the indirect flight muscle. Mol Biol Cell, 10(1):135–49, 1999.

23. H. Rommelaere, D. Waterschoot, K. Neirynck, J. Vandekerckhove, and C. Ampe. A method for rapidly screening functionality of actin mutants and tagged actins. Biol Proced Online, 6: 235–249, 2004.

24. A. Simiczyjew, A. J. Mazur, E. Dratkiewicz, and D. Nowak. Involvement of beta- and gamma-actin isoforms in actin cytoskeleton organization and migration abilities of bleb-forming human colon cancer cells. PLoS One, 12(3):e0173709, 2017.

25. A. Simiczyjew, A. J. Mazur, A. Popow-Wozniak, M. Malicka-Blaszkiewicz, and D. Nowak. Effect of overexpression of beta- and gamma-actin isoforms on actin cytoskeleton organization and migration of human colon cancer cells. Histochem Cell Biol, 142(3):307–22, 2014.

26. J. Q. Wu and T. D. Pollard. Counting cytokinesis proteins globally and locally in fission yeast. Science, 310(5746):310–4, 2005.

27. B. Roberts, A. Haupt, A. Tucker, T. Grancharova, J. Arakaki, M. A. Fuqua, A. Nelson, C. Hookway, S. A. Ludmann, I. A. Mueller, R. Yang, R. Horwitz, S. M. Rafelski, and R. N. Gunawardane. Systematic gene tagging using crispr/cas9 in human stem cells to illuminate cell organization. Mol Biol Cell, 28(21):2854–2874, 2017.

28. M. Westphal, A. Jungbluth, M. Heidecker, B. Muhlbauer, C. Heizer, J. M. Schwartz, G. Marriott, and G. Gerisch. Microfilament dynamics during cell movement and chemotaxis monitored using a gfp-actin fusion protein. Curr Biol, 7(3):176–83, 1997.

29. A. Drazic, H. Aksnes, M. Marie, M. Boczkowska, S. Varland, E. Timmerman, H. Foyn, N. Glomnes, G. Rebowski, F. Impens, K. Gevaert, R. Dominguez, and T. Arnesen. Naa80 is actin’s n-terminal acetyltransferase and regulates cytoskeleton assembly and cell motility. Proc Natl Acad Sci U S A, 115(17):4399–4404, 2018.

30. M. Goris, R. S. Magin, H. Foyn, L. M. Myklebust, S. Varland, R. Ree, A. Drazic, P. Bhambra, S. I. Stove, M. Baumann, B. E. Haug, R. Marmorstein, and T. Arnesen. Structural determinants and cellular environment define processed actin as the sole substrate of the n-terminal acetyltransferase naa80. Proc Natl Acad Sci U S A, 115(17):4405–4410, 2018.

31. Q. Chen, S. Nag, and T. D. Pollard. Formins filter modified actin subunits during processive elongation. J Struct Biol, 177(1):32–9, 2012.

32. L. R. Otterbein, P. Graceffa, and R. Dominguez. The crystal structure of uncomplexed actin in the adp state. Science, 293(5530):708–11, 2001.

33. M. Boczkowska, Z. Yurtsever, G. Rebowski, M. J. Eck, and R. Dominguez. Crystal structure of leiomodin 2 in complex with actin: A structural and functional reexamination. Biophys J, 113(4):889–899, 2017.

34. A. Einhauer and A. Jungbauer. Affinity of the monoclonal antibody m1 directed against the flag peptide. J Chromatogr A, 921(1):25–30, 2001.

35. H. Gotzke, M. Kilisch, M. Martinez-Carranza, S. Sograte-Idrissi, A. Rajavel, T. Schlichthaerle, N. Engels, R. Jungmann, P. Stenmark, F. Opazo, and S. Frey. The alfa-tag is a highly versatile tool for nanobody-based bioscience applications. Nat Commun, 10(1): 4403, 2019.

36. D. Agudelo, A. Duringer, L. Bozoyan, C. C. Huard, S. Carter, J. Loehr, D. Synodinou, M. Drouin, J. Salsman, G. Dellaire, J. Laganiere, and Y. Doyon. Marker-free coselection for crispr-driven genome editing in human cells. Nat Methods, 14(6):615–620, 2017.

37. P. Vedula and A. Kashina. The makings of the ’actin code’: regulation of actin’s biological function at the amino acid and nucleotide level. J Cell Sci, 131(9), 2018.

38. K. van den Dries, L. Nahidiazar, J. A. Slotman, M. B. M. Meddens, E. Pandzic, B. Joosten, M. Ansems, J. Schouwstra, A. Meijer, R. Steen, M. Wijers, J. Fransen, A. B. Houtsmuller, P. W. Wiseman, K. Jalink, and A. Cambi. Modular actin nano-architecture enables podosome protrusion and mechanosensing. Nat Commun, 10(1):5171, 2019.

39. D. Akhuli, A. Dhar, A. S. Viji, B. Bhojappa, and S. Palani. Aliby: Alfa nanobody-based toolkit for imaging and biochemistry in yeast. mSphere, 7(5):e0033322, 2022. ISSN 2379-5042 (Electronic) 2379-5042 (Linking). doi: 10.1128/msphere.00333-22.

40. D. Winter, A. V. Podtelejnikov, M. Mann, and R. Li. The complex containing actin-related proteins arp2 and arp3 is required for the motility and integrity of yeast actin patches. Curr Biol, 7(7):519–29, 1997. ISSN 0960-9822 (Print) 0960-9822 (Linking). doi: 10.1016/s0960-9822(06)00223-5.

41. A. E. Adams and J. R. Pringle. Relationship of actin and tubulin distribution to bud growth in wild-type and morphogenetic-mutant saccharomyces cerevisiae. J Cell Biol, 98(3):934–45, 1984. ISSN 0021-9525 (Print) 1540-8140 (Electronic) 0021-9525 (Linking). doi: 10.1083/jcb.98.3.934.

42. T. Splettstoesser, K. C. Holmes, F. Noe, and J. C. Smith. Structural modeling and molecular dynamics simulation of the actin filament. Proteins, 79(7):2033–43, 2011.

43. C. E. Schutt, J. C. Myslik, M. D. Rozycki, N. C. Goonesekere, and U. Lindberg. The structure of crystalline profilin-beta-actin. Nature, 365(6449):810–6, 1993. ISSN 0028-0836 (Print) 0028-0836 (Linking). doi: 10.1038/365810a0.

44. P. J. McLaughlin, J. T. Gooch, H. G. Mannherz, and A. G. Weeds. Structure of gelsolin segment 1-actin complex and the mechanism of filament severing. Nature, 364(6439):685–92, 1993. ISSN 0028-0836 (Print) 0028-0836 (Linking). doi: 10.1038/364685a0.

45. R. Dominguez and K. C. Holmes. Actin structure and function. Annu Rev Biophys, 40: 169–86, 2011. doi: 10.1146/annurev-biophys-042910-155359.

46. J. von der Ecken, S. M. Heissler, S. Pathan-Chhatbar, D. J. Manstein, and S. Raunser. Cryo-em structure of a human cytoplasmic actomyosin complex at near-atomic resolution. Nature, 534(7609):724–8, 2016. ISSN 1476-4687 (Electronic) 0028-0836 (Linking). doi:10.1038/nature18295.

47. K. Tanaka, S. Takeda, K. Mitsuoka, T. Oda, C. Kimura-Sakiyama, Y. Maeda, and Narita. Structural basis for cofilin binding and actin filament disassembly. Nat Commun, 9(1):1860, 2018. ISSN 2041-1723 (Electronic) 2041-1723 (Linking). doi: 10.1038/s41467-018-04290-w.

48. D. S. Kudryashov and E. Reisler. Atp and adp actin states. Biopolymers, 99(4):245–56, 2013.

49. H. Rommelaere, D. Waterschoot, K. Neirynck, J. Vandekerckhove, and C. Ampe. Structural plasticity of functional actin: pictures of actin binding protein and polymer interfaces. Structure, 11(10):1279–89, 2003.

50. Z. Wang, M. Grange, S. Pospich, T. Wagner, A. L. Kho, M. Gautel, and S. Raunser. Structures from intact myofibrils reveal mechanism of thin filament regulation through nebulin. Science, 375(6582):eabn1934, 2022. ISSN 1095-9203 (Electronic) 0036-8075 (Linking). doi: 10.1126/science.abn1934.

51. M. Boiero Sanders, C. P. Toret, A. Guillotin, A. Antkowiak, T. Vannier, R. C. Robinson, and Michelot. Specialization of actin isoforms derived from the loss of key interactions with regulatory factors. EMBO J, 41(5):e107982, 2022. ISSN 1460-2075 (Electronic) 0261-4189 (Print) 0261-4189 (Linking). doi: 10.15252/embj.2021107982.

52. L. Blanchoin, R. Boujemaa-Paterski, C. Sykes, and J. Plastino. Actin dynamics, architecture, and mechanics in cell motility. Physiol Rev, 94(1):235–63, 2014. ISSN 1522-1210 (Electronic) 0031-9333 (Linking). doi: 10.1152/physrev.00018.2013.

53. T. Hatano, S. Alioto, E. Roscioli, S. Palani, S. T. Clarke, A. Kamnev, J. R. Hernandez-Fernaud, L. Sivashanmugam, Y. Lazo B. Chapa, A. M. E. Jones, R. C. Robinson, K. Sampath, M. Mishima, A. D. McAinsh, B. L. Goode, and M. K. Balasubramanian. Rapid production of pure recombinant actin isoforms in pichia pastoris. J Cell Sci, 131(8), 2018. ISSN 1477-9137 (Electronic) 0021-9533 (Print) 0021-9533 (Linking). doi: 10.1242/jcs.213827.

54. T. Hatano, L. Sivashanmugam, A. Suchenko, H. Hussain, and M. K. Balasubramanian. Pick-ya actin - a method to purify actin isoforms with bespoke key post-translational modifications. J Cell Sci, 133(2), 2020. ISSN 1477-9137 (Electronic) 0021-9533 (Linking). doi: 10.1242/jcs.241406.

55. S. S. Lam, J. D. Martell, K. J. Kamer, T. J. Deerinck, M. H. Ellisman, V. K. Mootha, and Y. Ting. Directed evolution of apex2 for electron microscopy and proximity labeling. Nat Methods, 12(1):51–4, 2015. ISSN 1548-7105 (Electronic) 1548-7091 (Linking). doi: 10.1038/nmeth.3179.

56. T. C. Branon, J. A. Bosch, A. D. Sanchez, N. D. Udeshi, T. Svinkina, S. A. Carr, J. L. Feldman, N. Perrimon, and A. Y. Ting. Efficient proximity labeling in living cells and organisms with turboid. Nat Biotechnol, 36(9):880–887, 2018. ISSN 1546-1696 (Electronic) 1087-0156 (Linking). doi: 10.1038/nbt.4201.

57. F. Sherman. Getting started with yeast. Methods Enzymol, 194:3–21, 1991. ISSN 0076-6879 (Print) 0076-6879 (Linking). doi: 10.1016/0076-6879(91)94004-v. Sherman, F eng GM12702/GM/NIGMS NIH HHS/ Research Support, U.S. Gov’t, P.H.S. 1991/01/01 Methods Enzymol. 1991; 194:3–21. doi: 10.1016/0076-6879(91)94004-v.

58. L. F. Pemberton. Preparation of yeast cells for live-cell imaging and indirect immunofluorescence. Methods Mol Biol, 1205:79–90, 2014. ISSN 1940-6029 (Electronic) 1064-3745 (Linking). doi: 10.1007/978-1-4939-1363-3_6.

59. S. G. McInally, J. Kondev, and B. L. Goode. Scaling of subcellular actin structures with cell length through decelerated growth. Elife, 10, 2021. ISSN 2050-084X (Electronic) 2050-084X (Linking). doi: 10.7554/eLife.68424.

60. S. G. McInally, J. Kondev, and B. L. Goode. Quantitative analysis of actin cable length in yeast. Bio Protoc, 12(9):e4402, 2022. ISSN 2331-8325 (Electronic) 2331-8325 (Linking). doi: 10.21769/BioProtoc.4402.

61. A. Suarez-Arnedo, F. Torres Figueroa, C. Clavijo, P. Arbelaez, J. C. Cruz, and C. Munoz-Camargo. An image j plugin for the high throughput image analysis of in vitro scratch wound healing assays. PLoS One, 15(7):e0232565, 2020. ISSN 1932-6203 (Electronic) 1932-6203 (Linking). doi: 10.1371/journal.pone.0232565.

62. C. A. Schneider, W. S. Rasband, and K. W. Eliceiri. Nih image to imagej: 25 years of image analysis. Nat Methods, 9(7):671–5, 2012. ISSN 1548-7105 (Electronic) 1548-7091 (Linking). doi: 10.1038/nmeth.2089.

63. J. Goedhart. Superplotsofdata-a web app for the transparent display and quantitative comparison of continuous data from different conditions. Mol Biol Cell, 32(6):470–474, 2021. ISSN 1939-4586 (Electronic) 1059-1524 (Linking). doi: 10.1091/mbc.E20-09-0583.

