## Supplemental Figures and Tables for "IntAct: a non-disruptive internal tagging strategy to study actin isoform organization and function"

### Van Zwam et al. Supplementary Figure 1

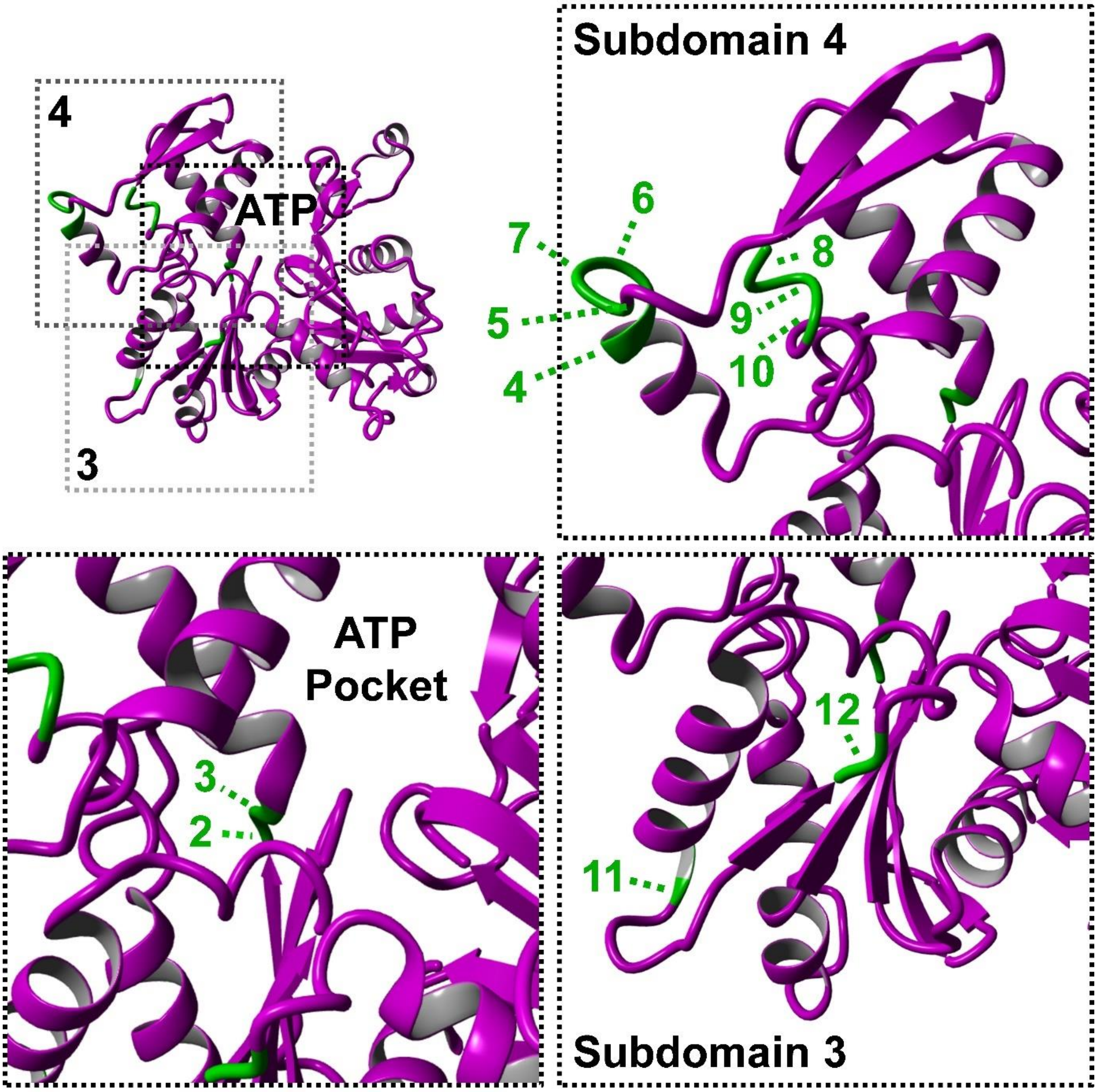

**Supplementary Figure 1 | Crystal structure of the actin domains and the position of each residue pair used in the screening.** Crystal structure of uncomplexed globular actin (magenta ribbon, PBD accession number: 1J6Z[32]) indicating subdomain 3, subdomain 4 and the ATP pocket. Zooms show each domain and ATP pocket and their associated position for each distinct residue pair.

### Van Zwam et al. Supplementary Figure 2

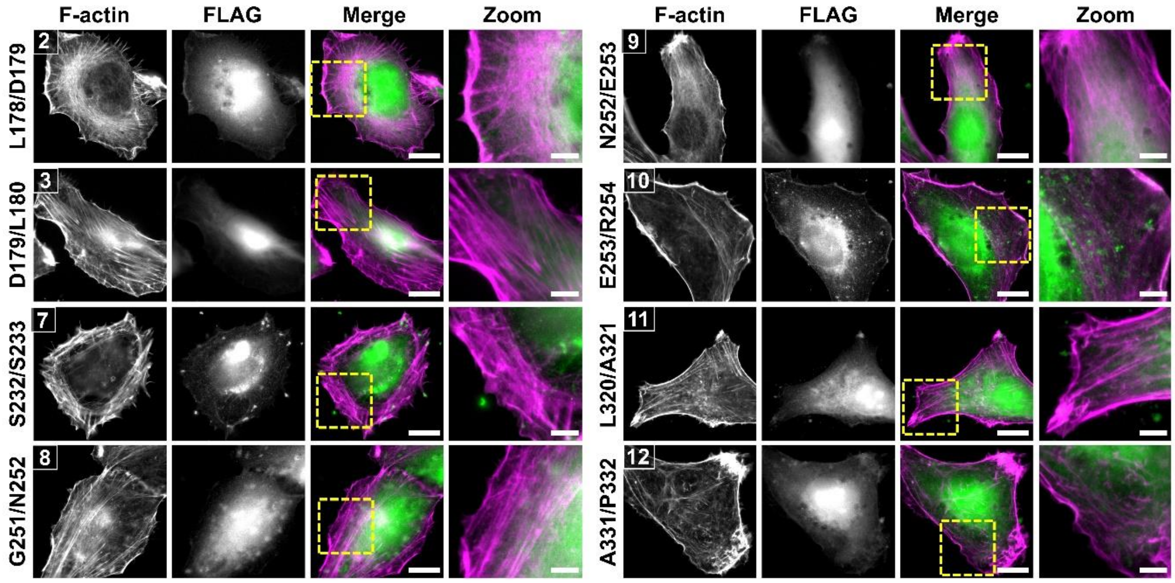

**Supplementary Figure 2 | Identification of actin T229/A230 as a permissive target site for epitope tag integration.** Representative widefield immunofluorescence images of F-actin (magenta) and FLAG (green) in HT1080 cells that overexpress the tagged β-actin variants. Shown are the eight internally tagged variants that are not depicted in **Fig. 1B**. Scale bar: 15 µm. Scale bar zoom: 5 µm.

### Van Zwam et al. Supplementary Figure 3

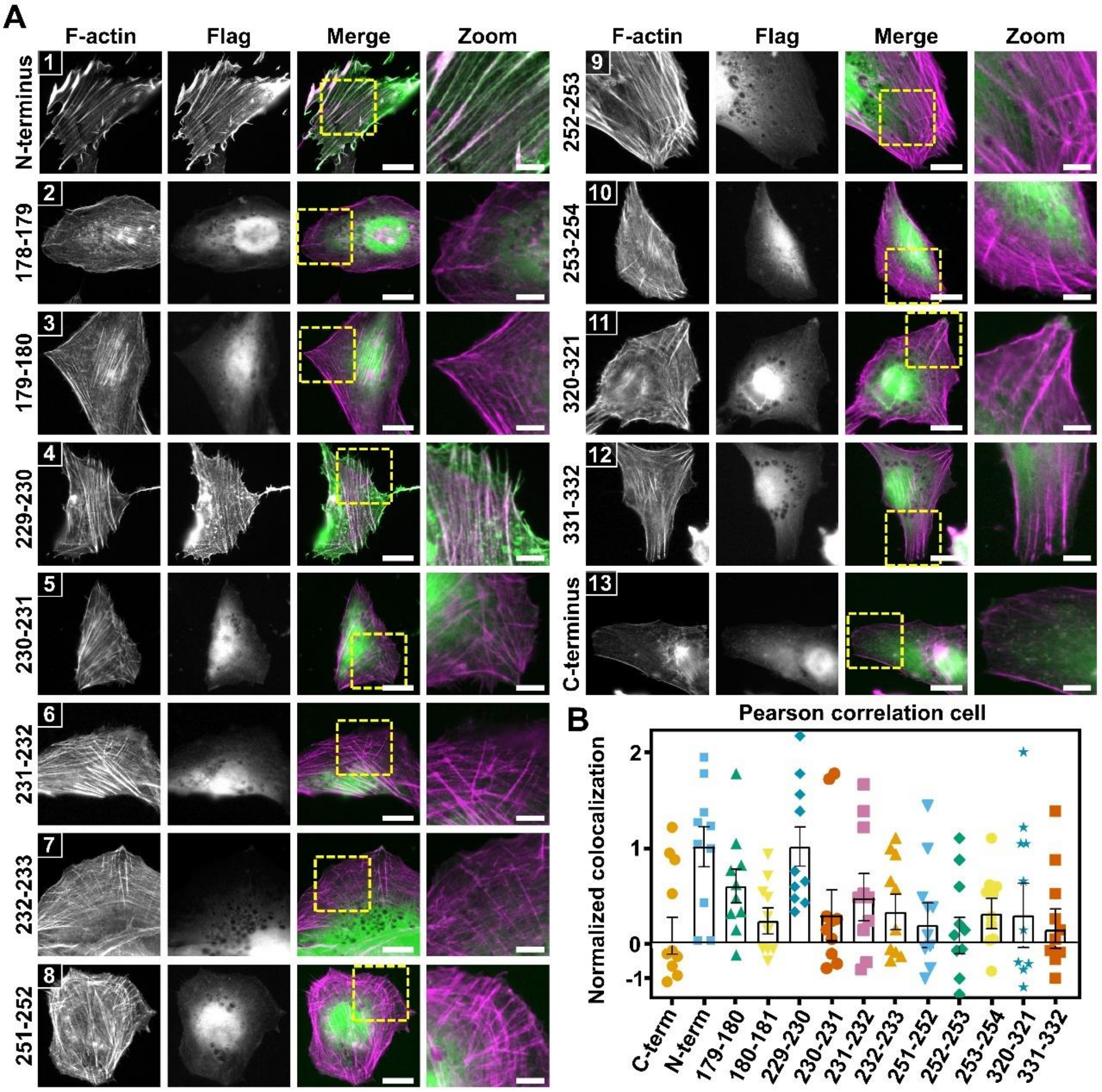

**Supplementary Figure 3 | Identification of actin T229/A230 as a permissive target site for epitope tag integration.** (**A**) Representative widefield immunofluorescence images of F-actin (magenta) and FLAG (green) in RPE1 cells that overexpress the tagged β-actin variants. Shown are eleven internally tagged variants and the N- and C-terminally tagged β-actin. Scale bar: 15 µm. Scale bar zoom: 5 µm. (**B**) Colocalization analysis of the microscopy results in **A** showing the normalized Pearson's correlation coefficient for each of the actin variants. Individual data points indicate single cells and in total, at least 10 cells from 2 independent experiments were included in the analysis. Bars represent the mean value, and error bars represent standard error of mean (SEM).

### Van Zwam et al. Supplementary Figure 4

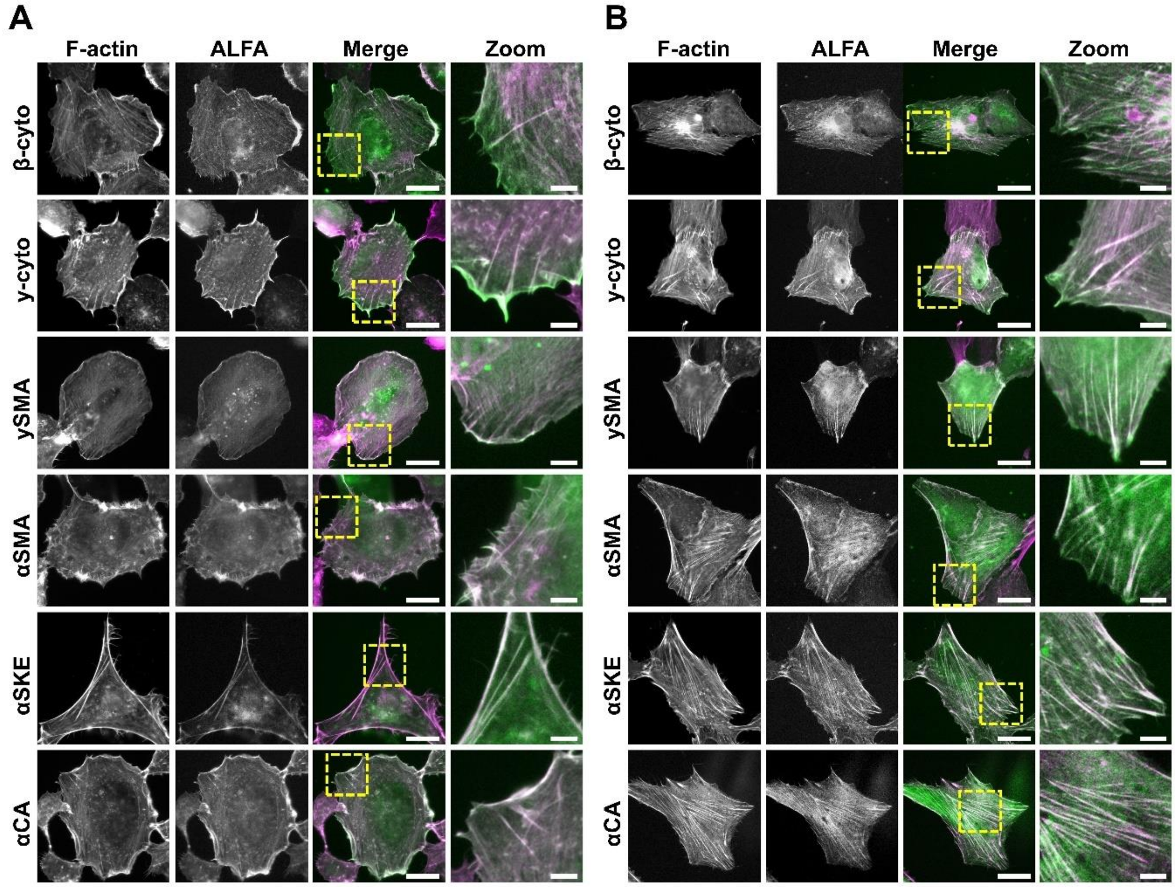

**Supplementary Figure 4 | Position T229/A230 is a permissive site for epitope integration in all six human actin isoforms.** (**A**) Representative widefield immunofluorescence images of HT1080 with F-actin (magenta) and ALFA tag (green) in HT1080 cells that overexpress the ALFA tag in position T229/A230 in all six human actin isoforms. Scale bar: 15 µm. Scale bar zoom: 5 µm. (**B**) Representative widefield immunofluorescence images of RPE1 with F-actin (magenta) and ALFA tag (green) in HT1080 cells that overexpress the ALFA tag in position T229/A230 in all six human actin isoforms. Scale bar: 15 µm. Scale bar zoom: 5 µm.

### Van Zwam et al. Supplementary Figure 5

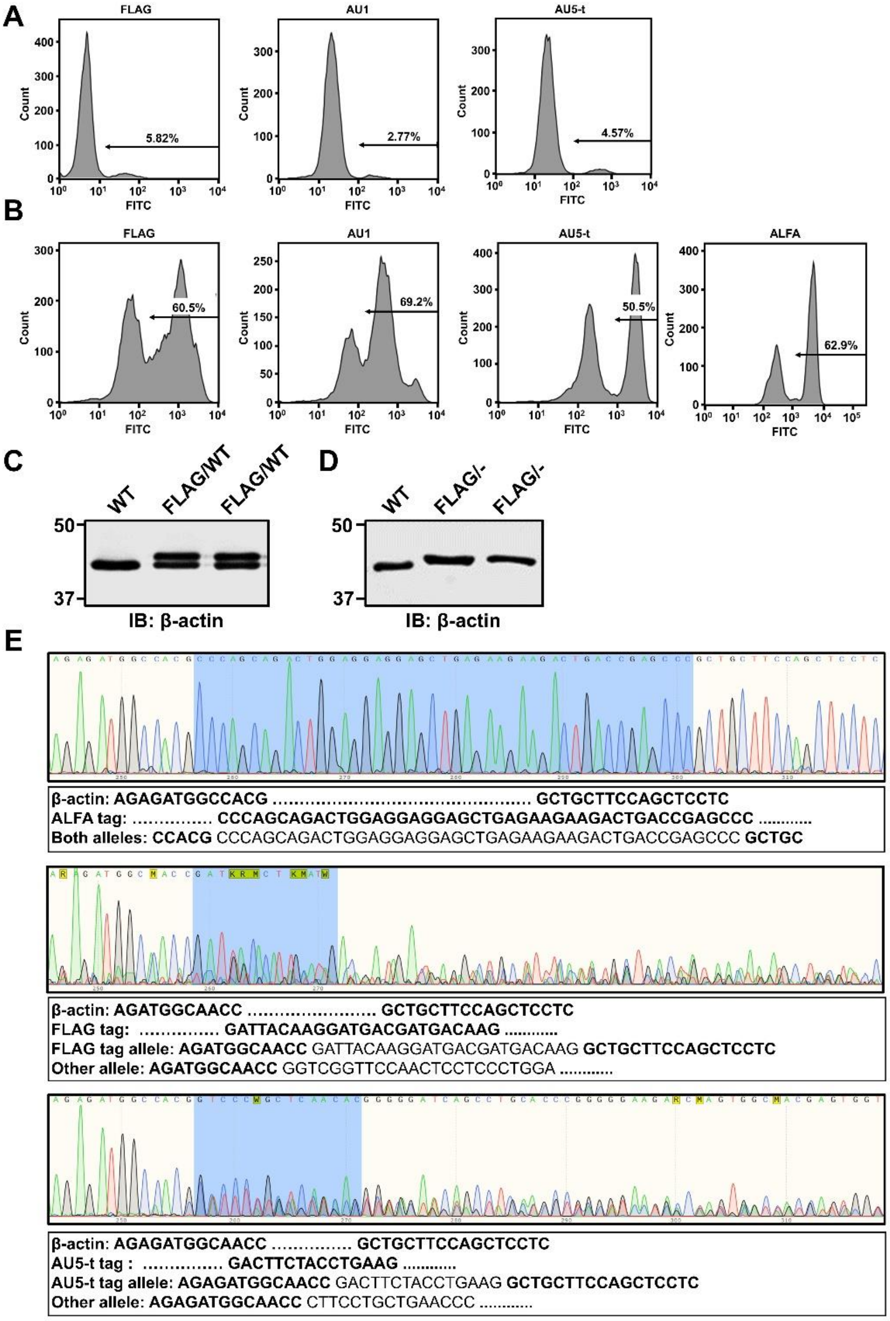

**Supplementary Figure 5 | Flow cytometry, western blot and Sanger sequencing data of cells having a CRISPR/Cas9-mediated knock-in of FLAG, AU1, AU5 or ALFA tag in β-actin.** (**A**) Flow cytometry data of cells having a CRISPR/Cas9-mediated knock-in of FLAG, AU1 or AU5. Pool of cells were stained for their appropriate tag. (**B**) Flow cytometry data of cells having a CRISPR/Cas9-mediated knock-in of FLAG, AU1, AU5 or ALFA after selection with Ouabain. Pool of cells were stained for their appropriate tag. (**C**) Representative western blot of β-actin in parental HT1080 (WT) and 2 independent heterozygous FLAG-β-actin HT1080 clones (FLAG/WT). (**D**) Representative western blot of β-actin in parental HT1080 (WT) and 2 independent hemizygous FLAG-β-actin HT1080 clones (FLAG/-). (**E**) Sanger sequencing result of homozygous ALFA-β-actin, hemizygous FLAG-β-actin and hemizygous AU5-β-actin HT1080 cells. Highlighted in blue is the ALFA, FLAG or AU5-t sequence at position T229/A230 in β-actin. Alignment of β-actin sequence, the tag sequence in one or two alleles and the possible disrupted allele sequence.

### Van Zwam et al. Supplementary Figure 6

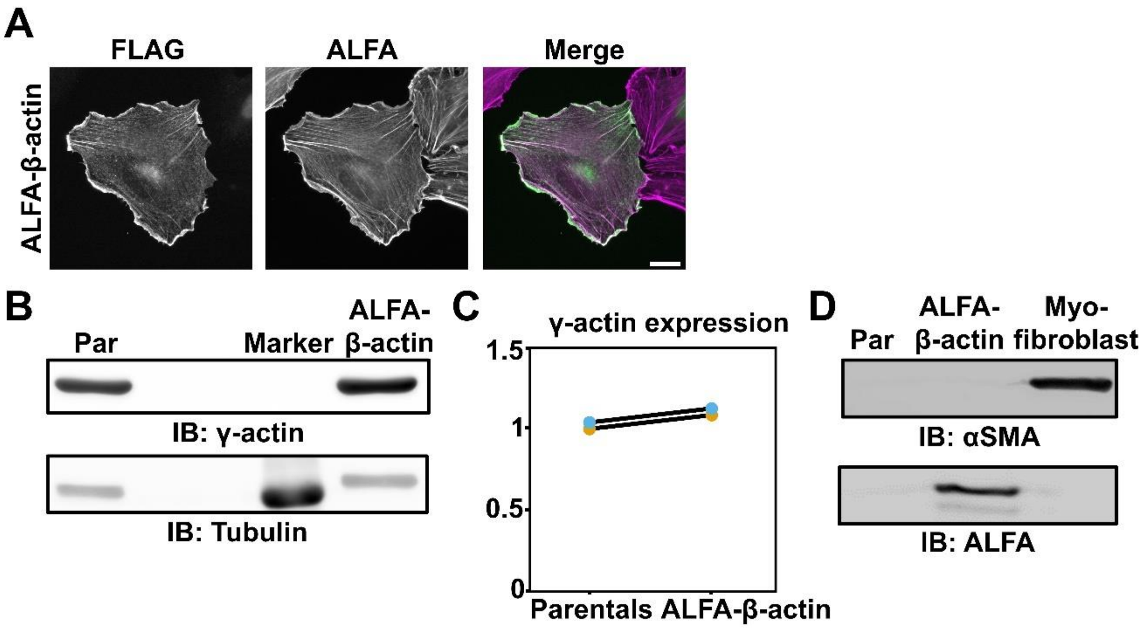

**Supplementary Figure 6 | Position T229/A230 is a permissive site for tagging both cytoplasmic isoactins and expression of γ-actin and αSMA is unaltered upon integration of ALFA-tag in β-actin.** (**A**) Representative widefield images from cells that have a CRISPR/Cas9-mediated knock-in of FLAG in γ-actin in ALFA-β-actin cells. Cells were labeled for ALFA (magenta) and FLAG (green) staining respectively for β-actin and γ-actin. Scale bar: 15 µm. (**B**) Representative western blot showing γ-actin expression in parental HT1080 (Par) and homozygous ALFA-β-actin HT1080 cells. Tubulin was used as a loading control. (**C**) Quantification of the γ-actin expression in ALFA-β-actin and parental HT1080 cells normalized to tubulin. (**D**) Representative western blot showing αSMA and ALFA tag expression in parental HT1080 (Par), homozygous ALFA-β-actin HT1080 cells and myofibroblast (positive control for αSMA).

### Van Zwam et al. Supplementary Figure 7

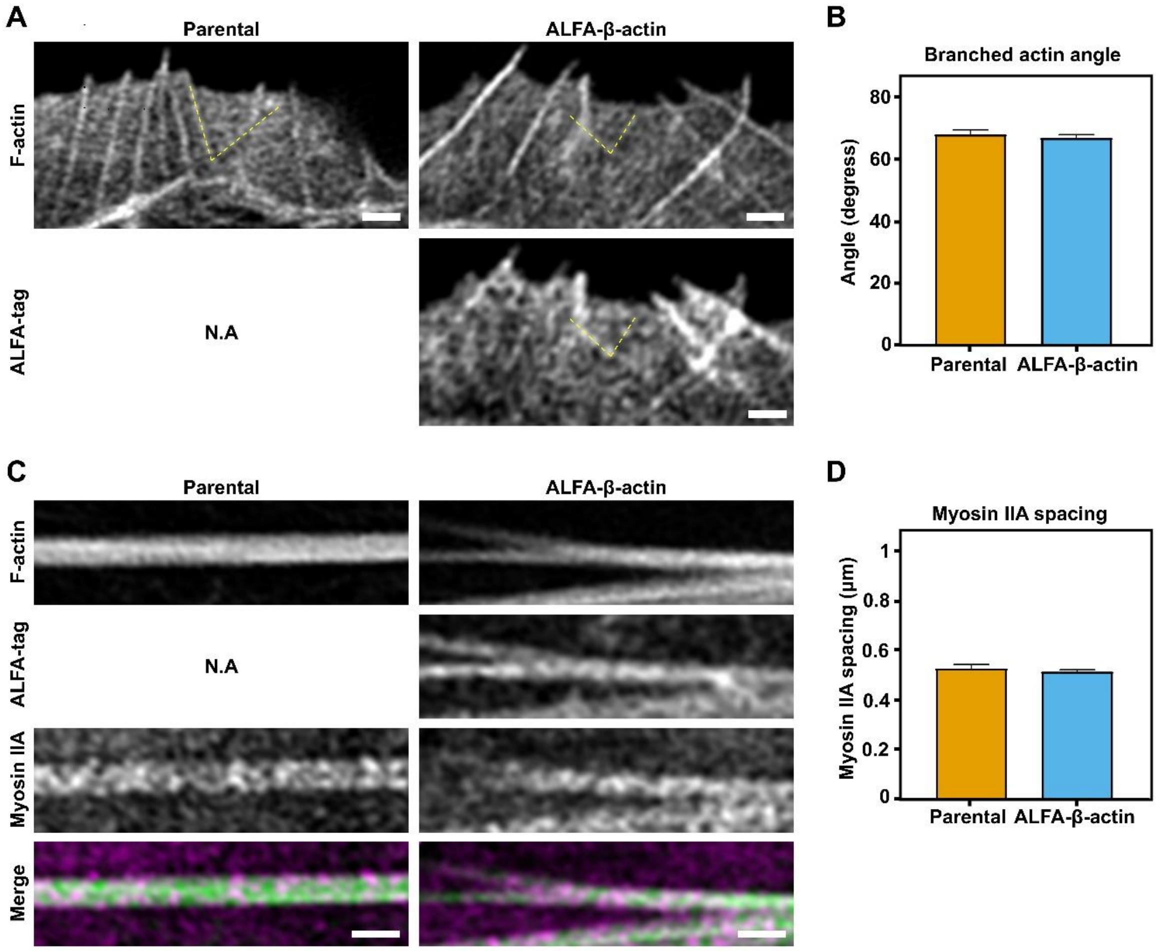

**Supplementary Figure 7 | Common actin-based structures have a similar architecture in ALFA-β-actin cells as compared to parental cells.** (**A**) Representative Airyscan images of lamellipodia from parental and ALFA-β-actin cells stained against F-actin and ALFA tag. Yellow dotted line follows the branched actin filaments. Scale bar: 1 µm. (**B**) Quantification of the angles of branched actin in **A**. Bars represent the mean value, and error bars represent standard error of mean (SEM) and at least 20 different branched actin angles were measured per condition. (**C**) Representative Airyscan images of stress fibers from parental and ALFA-β-actin cells stained against F-actin, ALFA tag and Myosin IIa. Scale bar: 1 µm. (**D**) Quantification of the myosin IIa spacing on stress fibers in **C**. Bars represent the mean value, and error bars represent standard error of mean (SEM) and in total the average myosin spacing of at least 30 stress fibers are included.

#### Van Zwam et al. Supplementary Figure 8

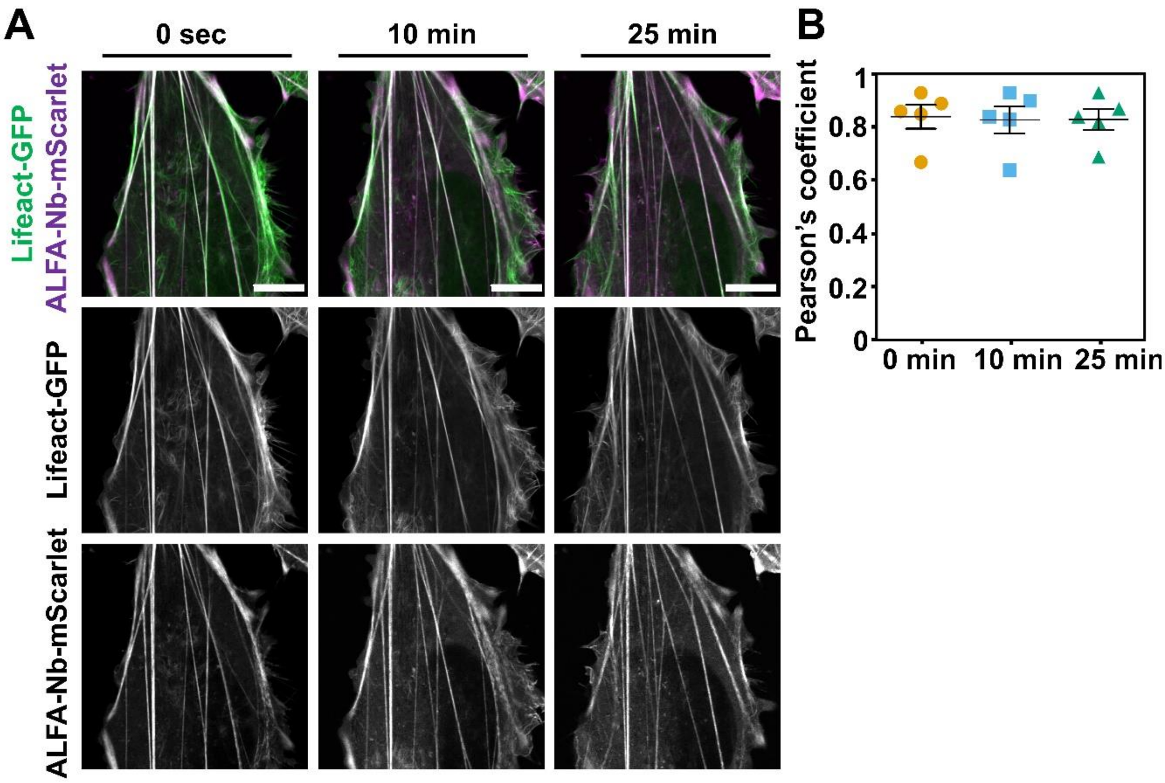

**Supplementary Figure 8 | ALFA-β-actin integrates properly into filaments in living cells.** (**A**) Representative Airyscan images of HT1080 cells expressing LifeAct-GFP (green) and ALFA-tag Nb-mScarlet (magenta) after 0 min, 10 min and 25 min. Full movie is available as **Suppl. Movie 1**. Scale bar: 10 µm. (**B**) Colocalization analysis of the microscopy results in **A** showing the Pearson's coefficient for each time point. Individual data points indicate single cells and in total, 5 different movies were included in the analysis. Bars represent the mean value, and error bars represent standard error of mean (SEM).

### Van Zwam et al. Supplementary Figure 9

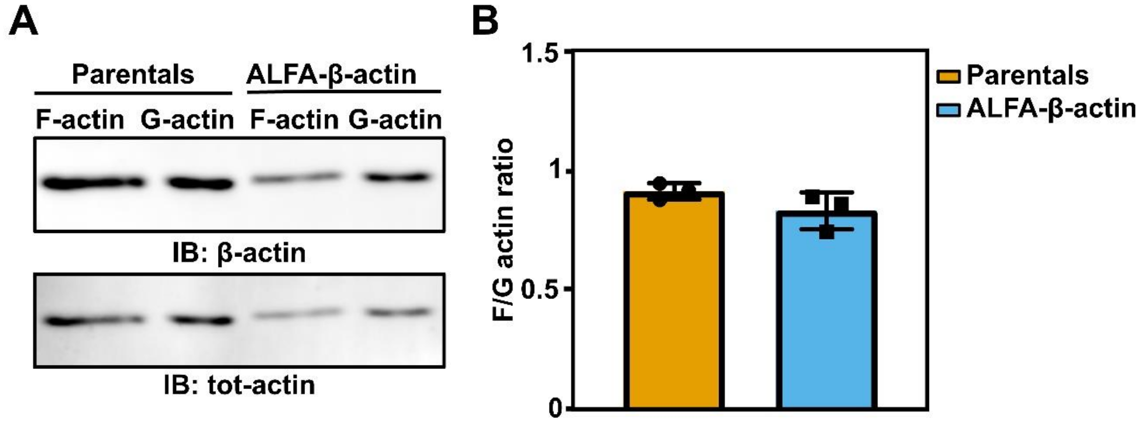

**Supplementary Figure 9 | F/G actin ratio unaltered upon tagging β-actin with ALFA tag at position T229/A230.** (**A**) Representative western blot of F-actin and G-actin fraction in parental HT1080 and homozygous ALFA-β-actin HT1080 cells. (**B**) Quantification of the F/G-actin ratio for β-actin from the western blots shown in **A**. Ratio were normalized against total actin. Bars represent the mean value, and error bars represent standard error of mean (SEM).

### Van Zwam et al. Supplementary Figure 10

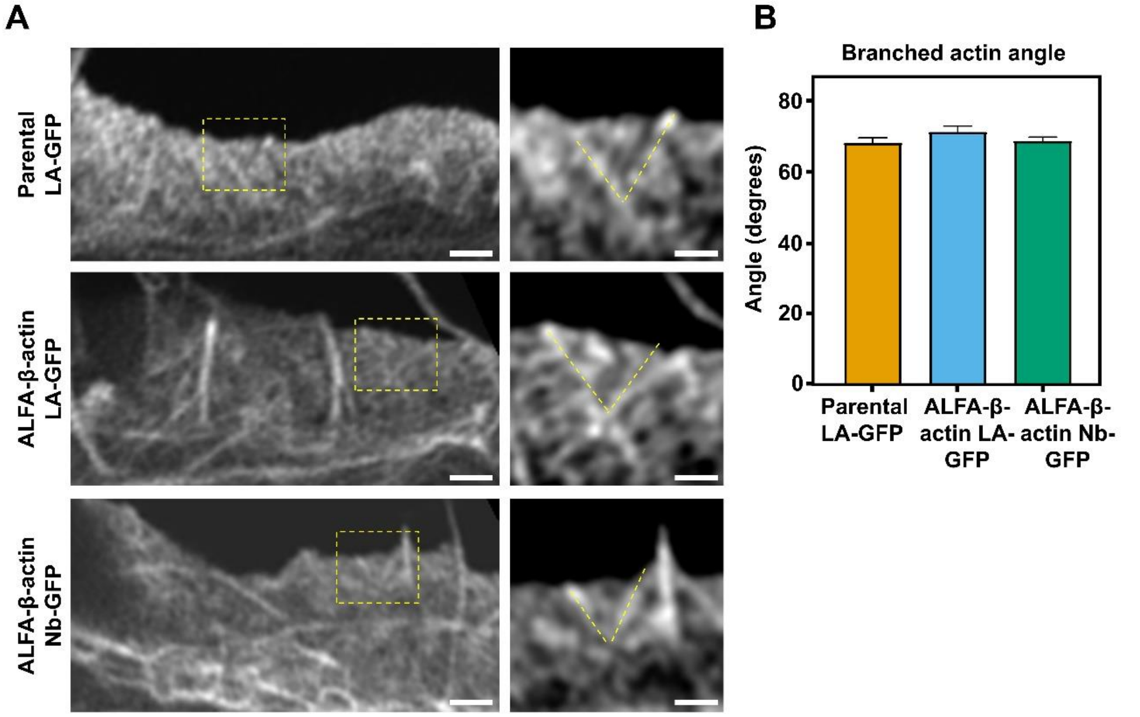

**Supplementary Figure 10 | Lamellipodia architecture is similar in ALFA-β-actin cells as compared to parental cells.** (**A**) Representative Airy scan images of lamellipodia from parental cells overexpressing LifeAct-GFP (LA-GFP), ALFA-β-actin cells overexpressing LifeAct-GFP (LA-GFP) or ALFA-β-actin cells overexpressing ALFA tag nanobody (Nb-GFP). Yellow square indicates zoom and yellow dotted line follows the branched actin filaments. Scale bar: 1 µm. Scale bar zoom: 0.5 µm. (**B**) Quantification of the angles of branched actin in **A**. Bars represent the mean value, and error bars represent standard error of mean (SEM) and at least 10 different branched actin angles were measured per condition.

### Van Zwam et al. Supplementary Figure 11

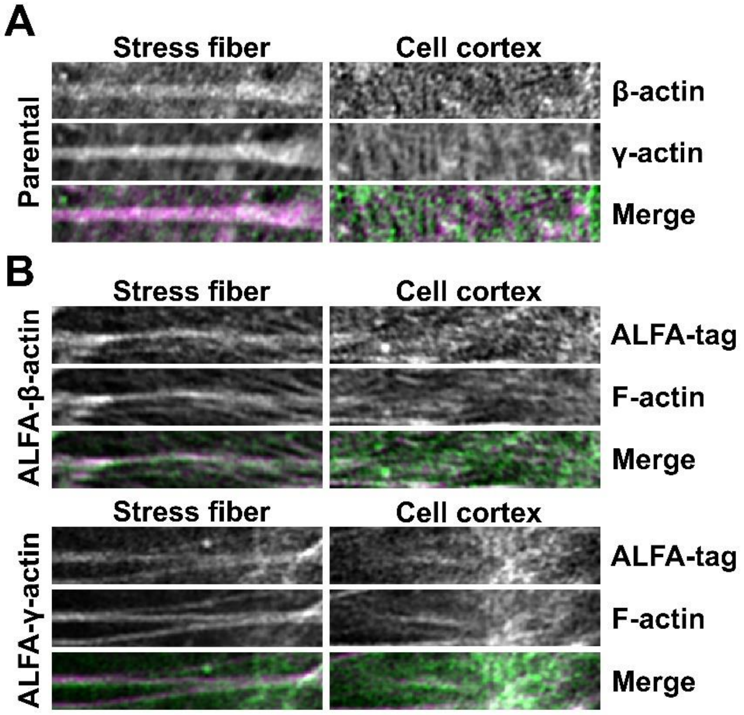

**Supplementary Figure 11 | IntAct β- and γ-actin recapitulate differential distribution of actin isoforms.** (**A**) Representative images of selected regions of interest of stress fibers (left column) and cell cortex (right column) in HT1080 parentals cells. Shown are β-actin (top row), γ-actin (second row) and merged (third row). (**B**) Representative images in β- and γ-actin IntAct cells of stress fibers (left column) and cell cortex (right column). Shown are ALFA tag (top row), F-actin (second row) and merged (third row).

### Van Zwam et al. Supplementary Figure 12

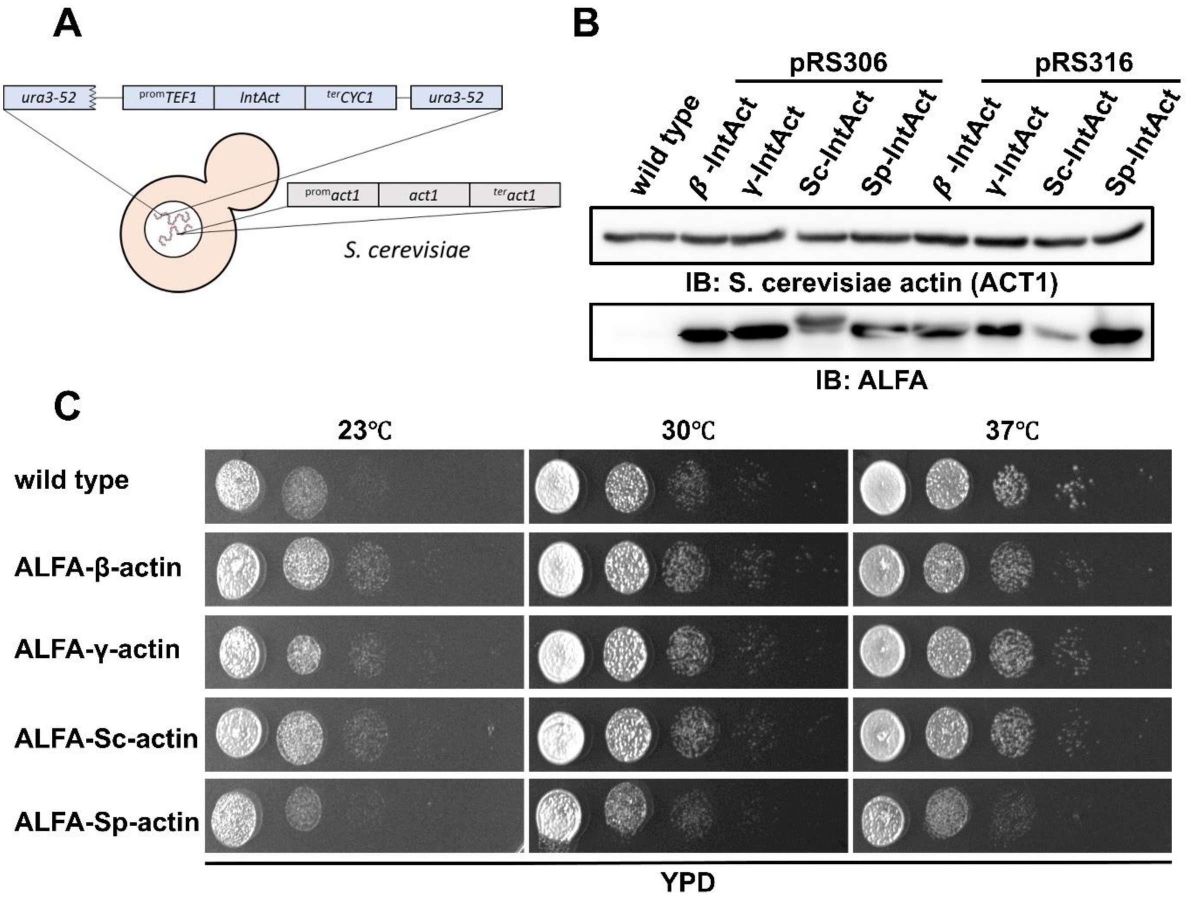

**Supplementary Figure 12 | Characterization of ALFA-tagged actins from yeast (Saccharomyces cerevisiae and Schizosaccharomyces pombe) and human (β- and γ-actin).** (**A**) Schematic overview showing an extra copy of IntAct actins (integrated at auxotrophic marker locus or present in plasmid) in addition to the native *S. cerevisiae act1* gene at its native locus. (**B**) Representative western blot showing expression of endogenous S. cerevisiae actin (ACT1) and β-IntAct, γ-IntAct, Sc-IntAct, Sp-IntAct expressed from either an integrating plasmid (pRS306) or centromeric plasmid (pRS316). (**C**) Spot test of strains expressing IntAct proteins with respect to wild type strain.

### Van Zwam et al. Supplementary Figure 13

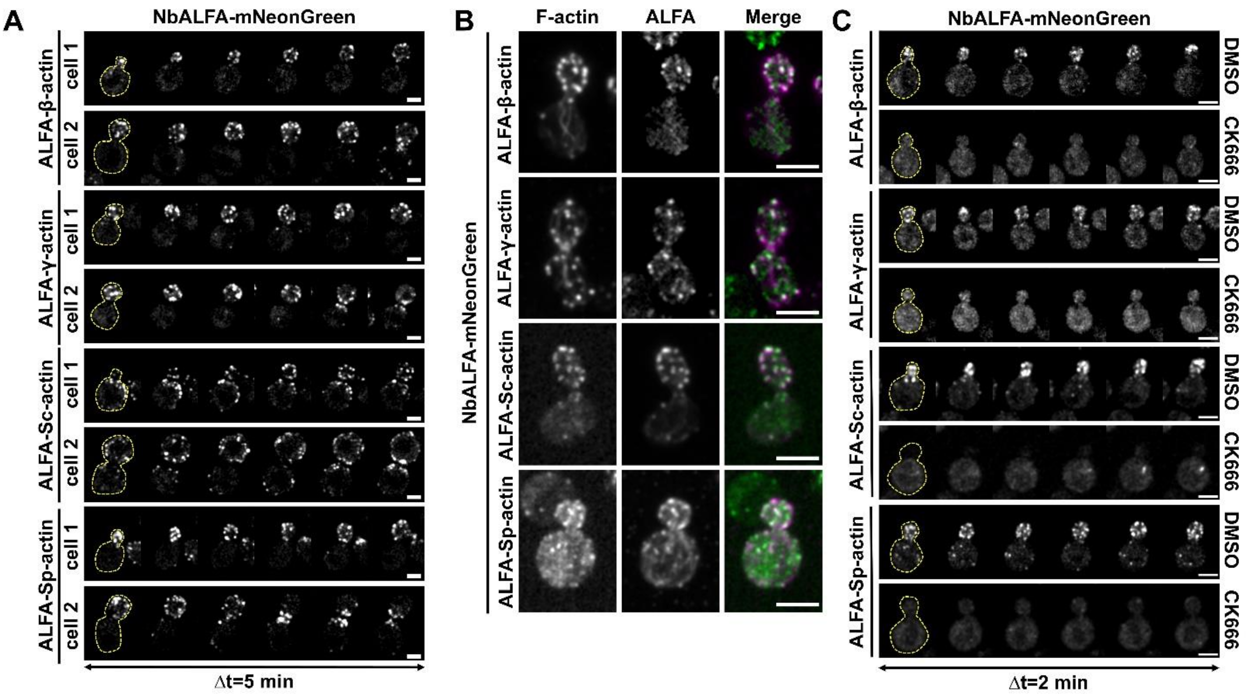

**Supplementary Figure 13 | IntAct actins localize to actin cortical patches in NbALFA-mNeonGreen expressing yeast.** (**A**) Representative confocal images of time-lapse imaging of NbALFA-mNeonGreen expressing yeast cells co-expressing β-IntAct, γ-IntAct, Sc-IntAct and Sp-IntAct. Yellow dashed line indicates the outline of the yeast cell. Scale bar: 3 µm. (**B**) Representative confocal images of NbALFA-mNeonGreen (green) expressing yeast cells co-expressing Sc-IntAct and Sp-IntAct. Stained for F-actin (magenta). Scale bar: 3 µm. (**C**) Representative montages of time-lapse imaging of NbALFA-mNeonGreen expressing yeast cells co-expressing: β-IntAct and γ-IntAct treated by either DMSO or CK666 (200µM). Yellow dashed line indicates the outline of the yeast cell. Scale bar: 3 µm.

### Van Zwam et al. Supplementary Figure 14

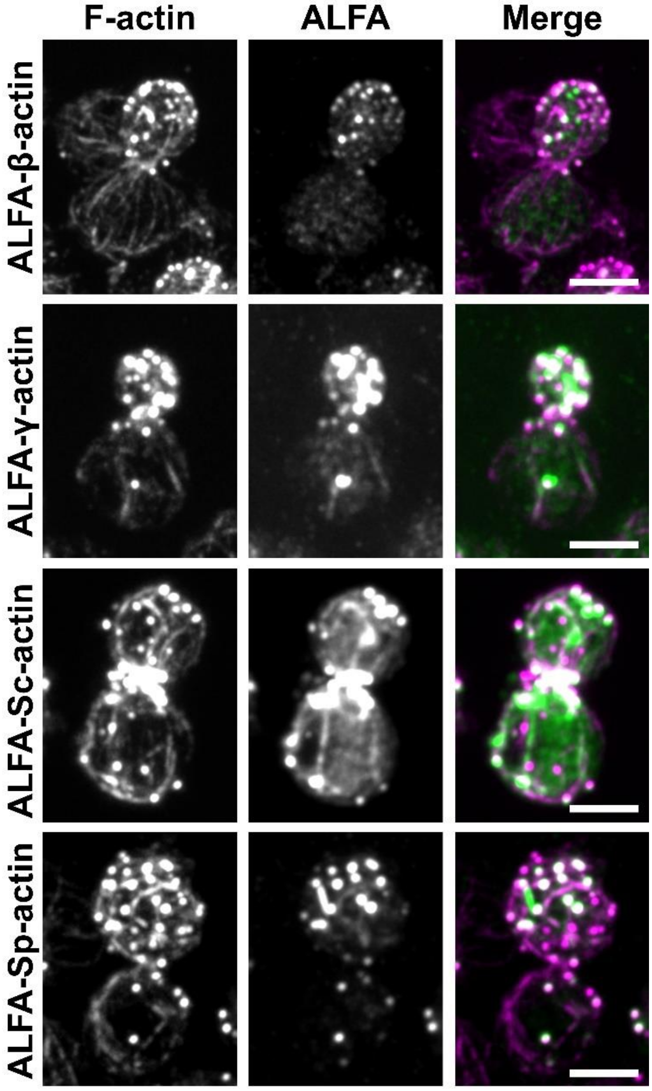

**Supplementary Figure 14 | IntAct actins localize to cortical actin patches and cytoplasmic actin filaments in wild type *S. cerevisiae.*** Representative montages of wild type yeast cells expressing β-IntAct, γ-IntAct Sc-IntAct and Sp-IntAct. Stained for F-actin (magenta) and ALFA tag nanobody (green). Scale bar: 3 µm.

### Van Zwam et al. Supplementary Table 1

| Strain Number | Genotype | Reference |
|---|---|---|
| YSP002 | ESM356 (MATa ura3-52 leu2Δ1 trp1Δ63 his3Δ200) *wild type* | Pereira and Schiebel, 2001[1] |
| YSP134 | YSP002 except leu2+ :pRS305-pTEF1- NbALFA-L-mNG-termCYC1 | Akhuli et. al., 2022[2] |
| YSP596 | YSP134 except ura3+ :pRS306-pTEF1-*β-IntAct*-termCYC1 | This study |
| YSP597 | YSP134 except ura3+ :pRS306-pTEF1-*γ-IntAct*-termCYC1 | This study |
| YSP600 | YSP134 except ura3+ :pRS306-pTEF1-*Sc-IntAct*-termCYC1 | This study |
| YSP601 | YSP134 except ura3+ :pRS306-pTEF1-*Sp-IntAct*-termCYC1 | This study |
| YSP641 | YSP002 except ura3+ :pRS316-pTEF1-*Sc-IntAct*-termCYC1 | This study |
| YSP642 | YSP002 except ura3+ :pRS316-pTEF1-*Sp-IntAct*-termCYC1 | This study |
| YSP643 | YSP002 except ura3+ :pRS316-pTEF1-*β-IntAct*-termCYC1 | This study |
| YSP644 | YSP002 except ura3+ :pRS316-pTEF1-*γ-IntAct*-termCYC1 | This study |

**Supplementary Table 1 |** List of Yeast Strains used in this study.

1. Pereira G, Schiebel E. The role of the yeast spindle pole body and the mammalian centrosome in regulating late mitotic events. Curr Opin Cell Biol. 2001 Dec 1;13(6):762–9.

2. Akhuli Dipayan, Dhar Anubhav, Viji Aileen Sara, Bhojappa Bindu, Palani Saravanan. ALIBY: ALFA Nanobody-Based Toolkit for Imaging and Biochemistry in Yeast. mSphere. 2022 Oct 3;7(5):e00333-22.

### Van Zwam et al. Supplementary Table 2

| Gene of interest: | Epitope insertion site: | Forward primer: (5'→3') | Reverse primer: (5'→3') |
|---|---|---|---|
| β-actin ORF | - | TATAGGGAGACCCAAGCTGGCTAGCATGGATGATGATATCGCCGCG | TTAGTGAACCGTCAGATCCGCTAGCATGGATGATGATATCGCCGCG |
| **Construct name:** | | | |
| Actb178/179 | L178^D179 | GATTACAAGGATGACGATGACAAGGACCTGGCTGGCCGGGACCTGAC | CTTGTCATCGTCATCCTTGTAATCCAGACGCAGGATGGCAT |
| Actb179/180 | D179^L180 | GATTACAAGGATGACGATGACAAGCTGGCTGGCCGGGACCTGACTGACTA | CTTGTCATCGTCATCCTTGTAATCGTCCAGACGCAGGATGGCAT |
| Actb229/230 | T229^A230 | GATTACAAGGATGACGATGACAAGGCTGCTTCCAGCTCCTCCCT | CTTGTCATCGTCATCCTTGTAATCCGTGGCCATCTCTTGCT |
| Actb230/231 | A230^A231 | GATTACAAGGATGACGATGACAAGGCTTCCAGCTCCTCCCTGGAGA | CTTGTCATCGTCATCCTTGTAATCAGCCGTGGCCATCTCTT |
| Actb231/232 | A231^S232 | GATTACAAGGATGACGATGACAAGTCCAGCTCCTCCCTGGAGAAGA | CTTGTCATCGTCATCCTTGTAATCAGCAGCCGTGGCCATCTCTT |
| Actb232/233 | S232^S233 | GATTACAAGGATGACGATGACAAGAGCTCCTCCCTGGAGAAGA | CTTGTCATCGTCATCCTTGTAATCGGAAGCAGCCGTGGCCATCTCTT |
| Actb251/252 | G251^N252 | GATTACAAGGATGACGATGACAAGAATGAGCGGTTCCGCTG | CTTGTCATCGTCATCCTTGTAATCGCCAATGGTGATGACCTGG |
| Actb252/253 | N252^E253 | GATTACAAGGATGACGATGACAAGGAGCGGTTCCGCTGCCCTGA | CTTGTCATCGTCATCCTTGTAATCATTGCCAATGGTGATGACCT |
| Actb253/254 | E253^R254 | GATTACAAGGATGACGATGACAAGCGGTTCCGCTGCCCTGAGGCA | CTTGTCATCGTCATCCTTGTAATCCTCATTGCCAATGGTGATGA |
| Actb320/321 | L320^A321 | GATTACAAGGATGACGATGACAAGGCACCCAGCACAATGAAGAT | CTTGTCATCGTCATCCTTGTAATCCAGGGCAGTGATCTCCTTCT |
| Actb331/332 | A331^P332 | GATTACAAGGATGACGATGACAAGCCTCCTGAGCGCAAGTACT | CTTGTCATCGTCATCCTTGTAATCAGCAATGATCTTGATCTTCATTGTG |
| Actb N-term | N-terminus | ATCGGCTAGCATGGATTACAAGGATGACGATGACAAGGATGATGATATCGCCGCG | CGATAAGCTTTTAGAAGCATTTGCGGTGGAC |
| Actb C-term | C-terminus | ATCGGCTAGCATGGATGATGATATCGCCGCG | CGTAAAGCTTTTACTTGTCATCGTCATCCTTGTAATCGAAGCATTTGCGGTGGACGAT |

**Supplementary Table 2 |** Forward and reverse primers used for the PCR fragments of the FLAG actin overexpression constructs.

### Van Zwam et al. Supplementary Table 3

| | |
|---|---|
| **gRNAs** | |
| *ACTB 229230* | 5’-GGAGGAGCTGGAAGCAGCCG-3’ |
| *ACTG1 229230* | 5’-AGAAGAGGAGGATGCGGCGG-3’ |
| *ATP1A1* | 5’-GAGTTCTGTAATTCAGCATA-3’ |
| **HDR templates** | |
| *ACTB* FLAG | 5’GAGAAGCTGTGCTACGTCGCCCTGGACTTCGAGCAAGAGATGGCAACCGATTACAAGGATGACGATGACAAGGCTGCTTCCAGCTCCTCCCTGGAGAAGAGCTACGAGCTGCCTGACGGC-3’ |
| *ACTB* AU5 | 5’ATTAAGGAGAAGCTGTGCTACGTCGCCCTGGACTTCGAGCAAGAGATGGCAACCGACTTCTACCTGAAGGCTGCTTCCAGCTCCTCCCTGGAGAAGAGCTACGAGCTGCCTGACGGCCAG-3’ |
| *ACTB* AU1 | 5’AAGGAGAAGCTGTGCTACGTCGCCCTGGACTTCGAGCAAGAGATGGCAACCGACACCTACAGATACATCGCTGCTTCCAGCTCCTCCCTGGAGAAGAGCTACGAGCTGCCTGACGGCCAG-3’ |
| *ACTB* ALFA | 5’GCTACGTCGCCCTGGACTTCGAGCAAGAGATGGCCACGCCCAGCAGACTGGAGGAGGAGCTGAGAAGAAGACTGACCGAGCCCGCTGCTTCCAGCTCCTCCCTGGAGAAGAGCTACGAGC-3’ |
| *ACTG1* ALFA | 5’GCTACGTCGCCCTGGACTTCGAGCAGGAGATGGCCACCCCCAGCAGACTGGAGGAGGAGCTGAGAAGAAGACTGACCGAGCCTGCCGCATCCTCCTCTTCTCTGGAGAAGAGCTACGAGC-3’ |
| *ATP1A1* mutations for ouabain resistance | 5’TGGAGCGATTCTTTGTTTCTTGGCTTATAGCATCAGAGCTGCTACAGAAGAGGAACCTCAAAACGATGACGTGAGTTCTGTAATTCAGCATATCGATTTGTAGTACACATCAGATATCTT -3’ |

**Supplementary Table 3 |** gRNAs and HDR templates used for insertion of FLAG-, AU5-, AU1- and ALFA-tags in position 229-230 of β-actin, the ALFA tag in γ-actin and HDR template for ouabain resistance. All oligos were inserted into the px330-hSpCas9 vector (Addgene, 42230) and the vector with the gRNA targeting *ATP1A1* was purchased (Addgene, 86611).

### Van Zwam et al. Supplementary Table 4

| Plasmid Number | Description | Reference |
|---|---|---|
| piSP1411 | pRS306-pTEF1-$\beta$-*IntAct*-termCYC1 | This study |
| piSP1412 | pRS306-pTEF1-$\gamma$-*IntAct*-termCYC1 | This study |
| piSP1413 | pRS306-pTEF1-*Sc*-*IntAct*-termCYC1 | This study |
| piSP1414 | pRS306-pTEF1-*Sp*-*IntAct*-termCYC1 | This study |
| piSP1415 | pRS316-pTEF1-$\beta$-*IntAct*-termCYC1 | This study |
| piSP1416 | pRS316-pTEF1-$\gamma$-*IntAct*-termCYC1 | This study |
| piSP1417 | pRS316-pTEF1-*Sc*-*IntAct*-termCYC1 | This study |
| piSP1418 | pRS316-pTEF1-*Sp*-*IntAct*-termCYC1 | This study |

**Supplementary Table 4 |** List of plasmid constructed and used in this study.

### Van Zwam et al. Supplementary Movie Legends

**Supplementary Video 1**
ALFA-tag colocalization with actin in HT1080 ALFA-β-actin cells transfected with LifeAct-GFP and Nanobody-mScarlet. Airyscan images were taken on a Leica LSM880 with 1 image per 15s. Playback speed: 15 fps, Scale bar: 10 µm.

**Supplementary Video 2**
Actin flow at the lamellipodia in HT1080 parental cells transfected with LifeAct-GFP. Airyscan images were taken on a Leica LSM880 with 1 image per 5s. Playback speed: 15 fps, Scale bar: 4 µm.

**Supplementary Video 3**
Actin flow at the lamellipodia in HT1080 ALFA-β-actin cells transfected with LifeAct-GFP. Airyscan images were taken on a Leica LSM880 with 1 image per 5s. Playback speed: 15 fps, Scale bar: 4 µm.

**Supplementary Video 3**
Actin flow at the lamellipodia in HT1080 ALFA-β-actin cells transfected with Nanobody-GFP. Airyscan images were taken on a Leica LSM880 with 1 image per 5s. Playback speed: 15 fps, Scale bar: 4 µm.
